# Optical mapping of ground reaction force dynamics in freely behaving *Drosophila melanogaster* larvae

**DOI:** 10.1101/2022.10.21.513016

**Authors:** Jonathan H. Booth, Andrew T. Meek, Nils M. Kronenberg, Stefan R. Pulver, Malte C. Gather

## Abstract

During locomotion, soft-bodied terrestrial animals solve complex control problems at substrate interfaces, but our understanding of how they achieve this without rigid components remains incomplete. Here, we develop new all-optical methods based on optical interference in a deformable substrate to measure ground reaction forces (GRFs) with micrometre and nanonewton precision in behaving *Drosophila* larvae. Combining this with a kinematic analysis of substrate interfacing features, we shed new light onto the biomechanical control of larval locomotion. Crawling in larvae measuring ∼1 mm in length involves an intricate pattern of cuticle sequestration and planting, producing GRFs of 1-7 µN. We show that larvae insert and expand denticulated, feet-like structures into substrates as they move, a process not previously observed in soft bodied animals. These ‘protopodia’ form dynamic anchors to compensate counteracting forces. Our work provides a framework for future biomechanics research in soft-bodied animals and promises to inspire improved soft-robot design.

## Introduction

Locomotion is a fundamental behaviour in the Animal Kingdom. There is great diversity in how it is accomplished, from the modification of torque angles in rigid bodied animals (1) to a diverse array of peristalses in limbed (2) and limbless soft-bodied animals (3). Key to these different strategies is one unifying characteristic: action against a substrate or fluid produces forces, thereby translating the body in space. In an aquatic environment, forces acting within fluids can be visualised via the waves of distortion they cause, thus facilitating the development of detailed theories of movement (4). In terrestrial settings, however, substrates are often rigid and therefore prevent direct visualisation of the ground reaction forces (GRFs) generated by animals.

Interactions with substrates have been extensively studied in animals with articulating skeletons (i.e. rigid bodied animals) due to the ability to calculate output forces from lever physics combined with measurements of joint-angles(1,5). However, much less is known about substrate interactions and GRFs in soft-bodied animals without rigid internal or external skeletons. These animals lack articulating joints upon which muscles act, ambiguating points through which the animal interacts with the substrate. However, they too must anchor a part of their body when another part is in motion to prevent net progression being impeded by an equal but opposite reaction force, i.e. their movements must obey Newton’s 3^rd^ law of motion (6). Furthermore, soft bodies pose a difficult control problem owing to their highly non-linear physical properties and virtually unlimited degrees of freedom. Movement over terrain therefore presents a unique challenge for soft animals. Dynamic anchoring has long since been postulated to be at the heart of soft-bodied locomotion (7), but understanding the mechanisms by which soft animals achieve this remains an open problem. Prior work on caterpillars (2,8–10), leeches (11,12) and *C. elegans* (13,14) provided key insights and have provided foundational observations for the inspiration of soft robot design; however, a lack of methods with sufficient spatiotemporal resolution for measuring GRFs in freely behaving animals has limited progress.

However, in the field of cellular mechanobiology, many new force measuring techniques have been developed which allow measurement of comparatively small forces from soft structures exhibiting low inertia (15–17) often with relatively high spatial-resolution. Early methods such as atomic force microscopy required the use of laser-entrained silicon probes to make contact with a cell of interest (15). This approach is problematic for studying animal behaviour due to the risk of the laser and probe influencing behaviour. Subsequently, techniques have been developed which allow indirect measurement of substrate interactions. One such approach is Traction Force Microscopy (TFM) in which the displacement of fluorescent markers suspended in a material with known mechanical properties relative to a 0-force reference allows for indirect measurement of horizontally aligned traction forces (17–19). This technique allows for probe-free measurement of forces, but has insufficient temporal resolution for the measurement of forces produced by many behaving animals, despite recent improvements (20). A second approach revolves around the use of micropillar arrays; in this technique, horizontally-aligned traction forces are measured by observing the deflection of pillars made of an elastic material with known mechanical properties. This approach can be limited in spatial resolution and introduces a non-physiological substrate that may influence animal behavior (21,22).

Recently we have introduced a technique named Elastic Resonator Interference Stress Microscopy (ERISM) which allows for the optical mapping of vertically aligned GRFs in the nanonewton range with micrometre precision by monitoring changes in local resonances of soft and deformable microcavities. This technique allows reference-free mapping of substrate interactions as well as calculation of vertically directed GRFs used in cell migration (23–25). Until recently, this technique was limited by its low temporal resolution (∼10s) making it unsuitable for use in recording substrate interaction during fast animal movements, but a very recent further development of ERISM known as wavelength alternating resonance pressure microscopy (WARP), has been demonstrated to achieve down to 10ms temporal resolution (26). Given ERISM and WARP allow for probe-free measurement of vertical ground reaction forces with high spatial and now temporal resolution, it becomes an attractive method for animal-scale mechanobiology.

In parallel, great strides have been made in understanding the neural and genetic underpinnings of locomotion in the *Drosophila* larva (27–31) a genetically tractable soft-bodied model organism (32). *Drosophila* larvae are segmentally organised peristaltic crawlers that move by generating waves of muscle contractions (3,31). Larvae have segmentally repeating bands comprised of 6 rows of actin trichomes (denticles) (33). The developmental and genetic origins of these structures have been extensively studied, but relatively little is known about how they are articulated during movement. While computational modelling and biomechanical measurements have provided an initial knowledgebase (34–36), data on biomechanical forces generated during substrate interactions in *Drosophila* larvae remain extremely limited (37,38). Development of methods for measuring GRFs in this model organism would enable fully integrated neurogenetic-biomechanical approaches to understanding soft-bodied movement and fulfil calls from the modelling community for more biomechanics data (39).

Here, we develop ERISM and WARP based approaches to measure GRFs exerted by freely behaving *Drosophila* larvae. We combine these measurements with kinematic tracking to explore how soft-bodied animals overcome fundamental biophysical challenges of moving over terrain. We find that, despite their legless appearance, *Drosophila* larvae interact with substrates by forming and articulating foot-like cuticular features (‘protopodia’) and cuticular papillae, which act as dynamic, travelling anchors. The use of ERISM-WARP provides a step-change in capability for understanding how soft-bodied animals interact with substrates and paves the way for a wider use of optical force measurement techniques in animal biomechanics and robotics research.

## Results

### Kinematic tracking of substrate interfacing features

As a first step in understanding how larvae interact with substrates, we confined 3^rd^ instar larvae to glass pipettes lined with soft agarose (0.1% w/v) (**Figure 1A**). This allowed us to laterally image the animals and the lateral edges of denticle rows at the substrate interface (**Figure 1B**) while animals crawled towards an appetitive odour source. Animals interact with the substrate by large, soft, segmentally repeating cuticular features that contain rows of denticles and to which we refer as ‘protopodia’ in the following. Protopodia in each segment engaged in ‘swing’ periods (moving, SwP) and ‘stance’ periods (planted on substrate, StP) as waves propagated through the body. During SwPs, protopodia detached from the substrate, with the posterior row of denticles moving to meet the anterior row of denticles, thereby inverting the cuticle and sequestering the whole protopodia into a travelling pocket (**Figure 1C**). When protopodia ended their SwP, they unfolded from the sequestration pocket and then protruded into the substrate during the StP.

**Figure 1.**
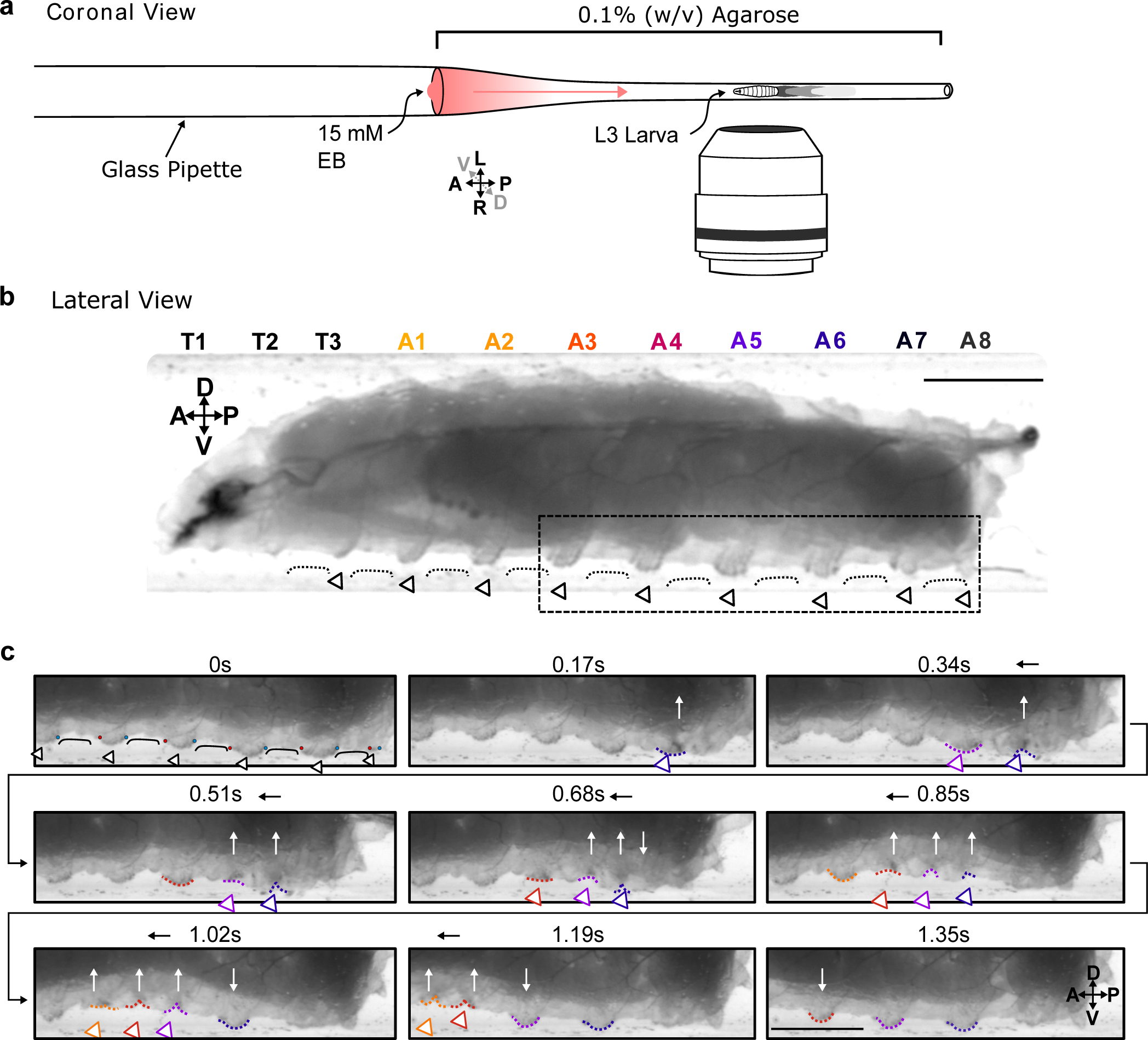
Protopodia protrusions in each segment are sequestered during swing phases of forwards locomotion. **a**, Schematic of setup for lateral imaging of larvae, using confinement in Pasteur pipette pre-filled with 0.1% (w/v) agarose. To encourage forward crawling, 10µl of 15mM ethyl butanoate (EB) was placed as attractive odour at the end of the pipette. **b**, Lateral brightfield image of 3^rd^ instar larva showing convex areas of denticle bands (open arrowheads) protruding into the substrate, interdigitated by concave areas of naked cuticle (black line) not interacting with the substrate. Scale bar=750µm. **c**, Time lapse of area marked by dotted box in **b** showing the swing periods and stance periods of protopodia (coloured open arrowheads and dotted lines) during a forward wave. Red and blue dots at 0s denote anterior and posterior rows of denticles, respectively. As the posterior-most denticle row moved to meet the anterior row of the band, the medial row detached from the substrate via invagination (white arrows). The invaginated pocket is then moved forward (black arrow) and subsequently replanted. This action repeats as the wave propagates. Scale bar=500µm. Images representative of three 3^rd^ instar larvae.

To further investigate the dynamics of protopodia placements, we performed detailed kinematic tracking of the morphometry of protopodia, denticle bands, and inter-protopodial spaces during peristaltic waves. By tracking the movement of defined points on bands relative to each other, we monitored intersegmental and intra-segmental movements during peristaltic waves (**Figure 2A**). In addition to moving relative to each other, denticle bands changed their shape during the sub-phases of a peristaltic wave. During forward waves (peristaltic contractions travelling in an anterograde direction), the anterior-most row of each denticle started to move after the corresponding posterior-most row (**Figure 2B**) and completed its movement after the posterior-most row stopped moving (**Figure 2C**), i.e. there was an anteroposterior (AP) latency for both Swing Initiation (SI) (when movement begins) and for Swing Termination (ST) (when movement ends). Such a ‘rolling’ progression pattern is analogous to the ‘heel-to-toe’ footfalls of limbed animals (40). To analyse this pattern further, we quantified the percentage of the wave duration spent in AP latency during SI and ST. For forwards waves, this relative latency was generally consistent across the denticle bands on large protrusive protopodia but less pronounced for the smaller and less protruding protopodia at the extreme posterior and anterior abdomen and the thorax (**Figure 2D**). In backwards waves, the heel-toe like latency was reversed, with anterior-led latencies observed in SI and posterior-led latencies observed in ST (**Supplementary Figure S1**).

**Figure 2.**
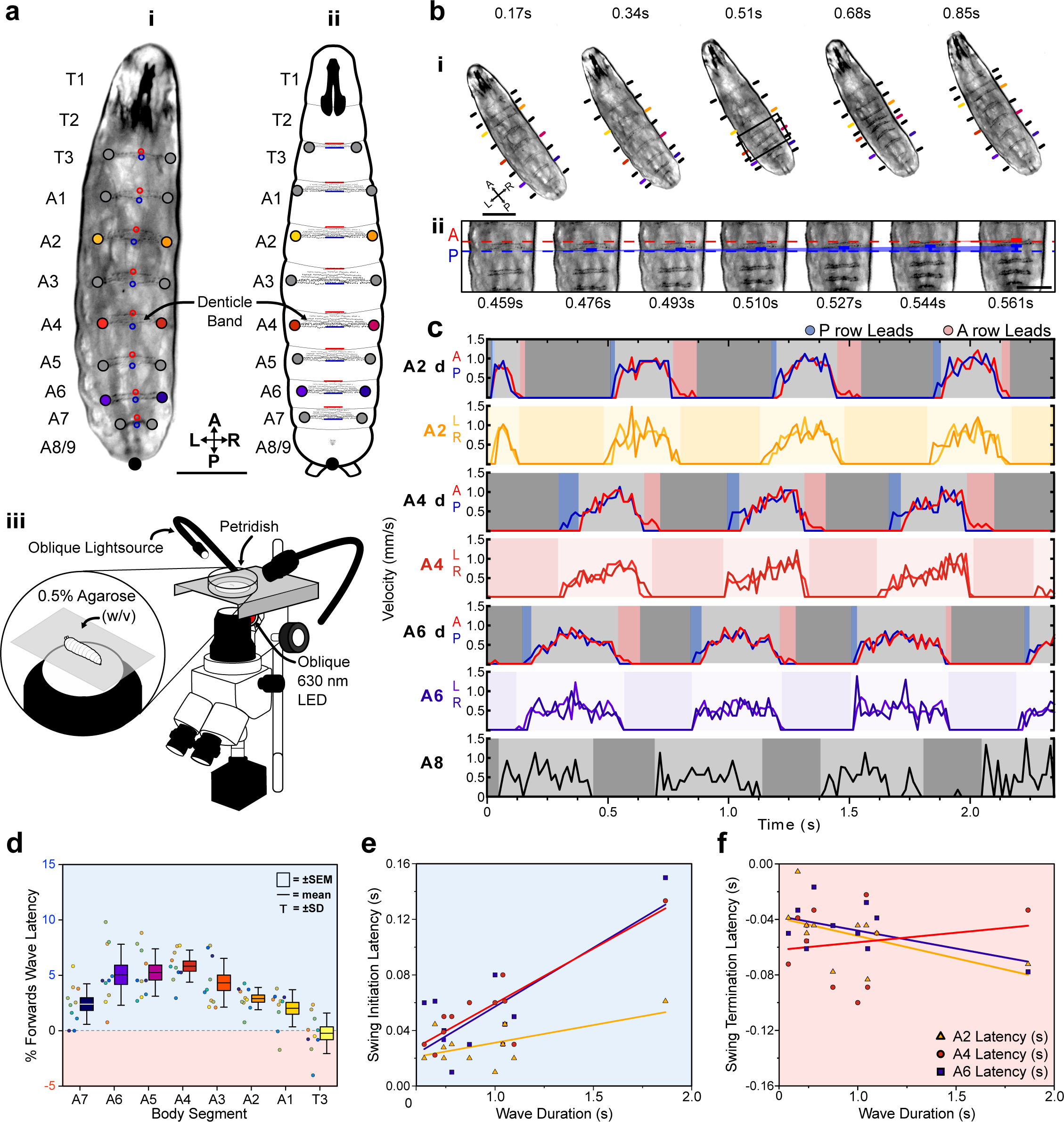
Protopodia kinematics follow ‘heel-toe’-like footfall dynamics. **a**, (**i**) Brightfield image and (**ii**) schematic of 2^nd^ instar larvae showing ventral side denticle belts which reside upon the protopodia and (**iii**) schematic of the imaging setup used for kinematic tracking. Scale bar=200µm. **b**, (**i**) As a forward wave travels through the animal, the distance between denticle bands decreases. Scale bar=200µm. (**ii**) At higher frame rate and magnification, changes in distance between the posterior and anterior most denticle rows are resolved. The posterior-most row (P, blue) initiates movement first and moves until nearly reaching the anterior-most row (A, red) at 0.544s, after which point, they move together (0.561s). Scale bar=100µm. **c**, Velocity of anterior– and posterior-most denticles rows (A2d A/P, A4d A/P, A6d A/P) and the left/right end of denticle bands (A2 L/R, A4 L/R, A6 L/R and A8 L/R) over three representative forward waves, showing how the strategy observed in **b** is maintained across body segments. Background colours indicate swing initiation (SI, blue), swing period (SwP, light grey), swing termination (ST, pink) and stance period (StP, dark grey). **d**, Forward wave latency for different animals and body segments. Positive values denote posterior row led latency. n=10 animals, 30 waves. **e**, SI-latency scales with wave duration in the posterior abdomen (A6: R^2^=0.61, purple; A4: R^2^=0.78, red) but less so for the anterior abdomen (A2: R^2^=0.35, yellow). n=12 animals with 3 latency periods per segment. **f**, ST-latencies do not scale with wave duration (A6: R^2^=0.26, A4: R^2^=0.26, A2: R^2^=0.03). n=12 animals with 3 latency periods per segment.

In summary, each segment-wise denticle action event is composed of four distinct periods: SI, SwP, ST, and StP. For forward waves and posterior segments, the latencies during the SI period are largely determined by wave duration (R^2^ range: 0.46-0.78, A7-A4) but this is less the case for anterior abdomen and thorax (R^2^ range: 0.12-0.35, A3-A1 and T3, **Figure 2E**). The magnitudes of ST-related latencies are not strongly strongly determined by wave duration (R^2^ range: 0.01-0.26, **Figure 2F**).

### Developing stress microscopy for *Drosophila*

Kinematic analysis of protopodia movements revealed a previously uncharacterised complexity in the dynamics of larval movement, but it cannot quantify the mechanical forces impacting the substrate and is therefore limited to making inferences regarding substrate interaction. To achieve quantitative observations, we therefore adapted ERISM-WARP (**Figure 3A**, **Supplementary Figure S2**) to map the vertically directed GRFs exerted by larvae rather than the forces exerted by single cells. First, we developed optical microcavities with mechanical stiffnesses in the range found in hydrogel substrates commonly used for studying *Drosophila* larval behaviour, i.e. Young’s modulus (E) of 10-30kPa (41–43). These microcavities consisted of two semi-transparent, flexible gold mirrors sandwiching a transparent polymer rubber that was made from a mixture of siloxanes with discrete Young’s moduli to adjust the resulting stiffness (44). The microcavities were characterised using atomic force microscopy (AFM) and the resulting force distance curves (**Figure 3B**) were fitted to a height-corrected Hertz Model to determine the Young’s modulus of each cavity (45). This procedure allowed us to fabricate microcavities with a wide range of well-defined Young’s moduli (**Figure 3C**, **Supplementary Table 1**).

**Figure 3.**
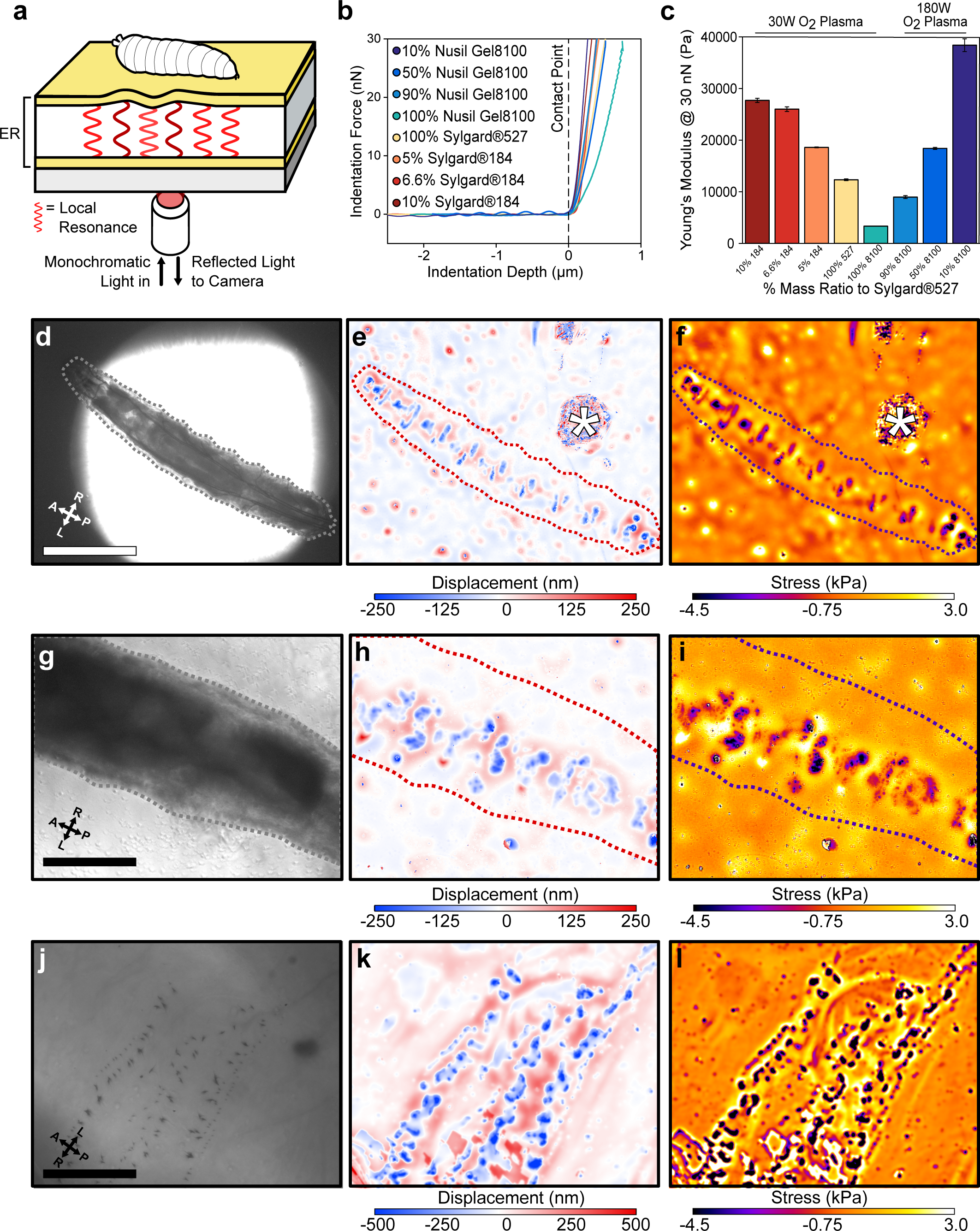
ERISM maps mechanical substrate interactions in *Drosophila* larvae. **a**, Schematic of setup for ERISM with *Drosophila* larva on an optical microcavity. Maps of local cavity deformation (displacement) due to indentation forces are generated by analysing cavity resonances. **b**, Force distance relationship measured by AFM and **c**, Mechanical stiffnesses (Young’s moduli) for microcavities produced by mixing different elastomers at different ratios and applying different plasma conditions. **d**, **g**, **j**, Brightfield images of anaesthetised 2^nd^ instar larvae recorded at low, medium, and high magnification. **e**, **h**, **k**, Corresponding maps of microcavity displacement. (* denotes contamination on cavity surface from handling the larva.) **f**, **i**, **l**, Corresponding maps of mechanical stress obtained by finite element analysis of displacement maps, showing the stress on the substrate due to passive interaction between larvae and substrate. Scale bar=500µm (**d**), 250µm (**g**) and 50µm (**j**). Images representative of 4 separate 2^nd^ instar larvae. Microcavities in **d**-**i** used 30W O_2_ 10% Sylgard®184 design, and **j**-**l** used a 30W O_2_ 5% Sylgard®184 design.

As an initial test, we placed cold-anaesthetised 2^nd^ instar larvae onto a microcavity (E=28kPa) and performed ERISM force mapping at different magnifications to record substrate indentations generated by larval body features (**Figure 3D-L**). Indentation maps were computed from the images of optical interference by pixelwise solving of the resonance condition with an optical model. Stress maps were then computed from the indentation maps via a finite element method (FEM) simulation of the stress distribution required to produce the observed indentation profile (Methods; the accuracy of our calculations was confirmed applying a known force with an AFM, **Supplementary Figure S5**). With this approach, we were able to resolve indentations from rows of denticle bands interdigitated by naked cuticle (**Figure 3G-I**). At higher magnification and when using slightly softer microcavities (E=19kPa), even indentations from individual denticles within these bands were resolved (**Figure 3J-L**). The median force exerted by individual denticles was 11.51nN (1.4nN-47.5nN; n=130 denticles) across a median area of 2.81µm (1.15-9.13µm; n=130 denticles).

### Videorate force mapping in freely behaving animals

Next, we moved to force mapping of freely behaving animals. First, we confirmed that ordinary larval behaviour is maintained on collagen-treated microcavity substrates (**Supplementary Figure S3**). We then adapted WARP (26) to image substrate interactions at high temporal resolution (**Supplementary Figure S4)**. For forward peristaltic waves, we observed posterior to anterior progressions of indentations into the cavity, corresponding to protopodial placements (**Figure 4A**). We also observed upward deflections of the substrate (i.e., increase in microcavity thickness, positive stress), associated with the displacement of elastomer because of Poisson’s ratio governing elastic materials (46). We also observed that the animals travel surrounded by a relatively large water droplet. During StP, protopodia displaced the substrate, and during SwP, protopodia local to the contraction were completely removed from the substrate while travelling to their new resting position.

**Figure 4.**
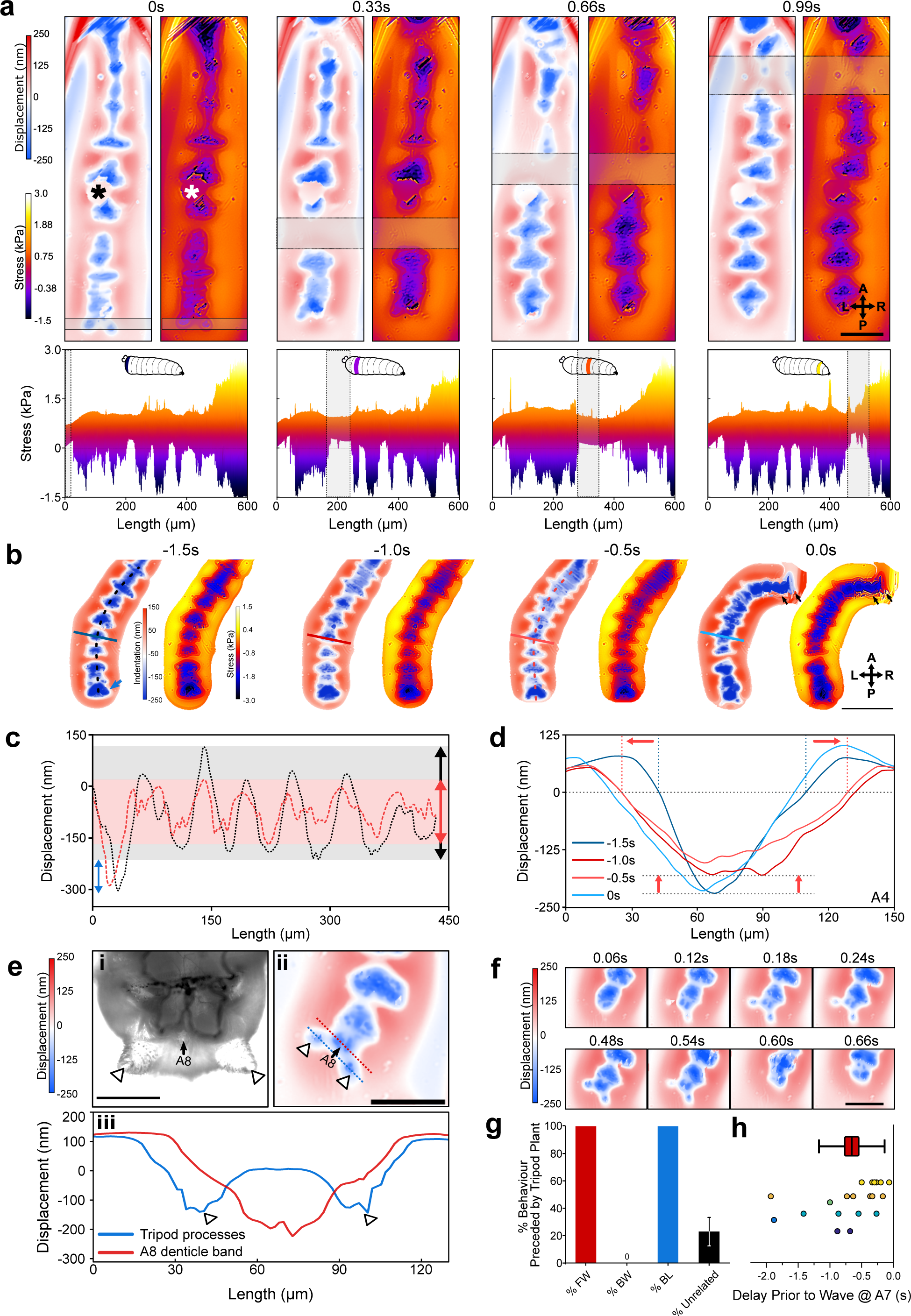
WARP imaging reveals dynamics of substrate interactions during larval movement. **a**, WARP image sequence of displacement and stress maps (top) for a freely behaving 2^nd^ instar larva during forward locomotion. (*denotes dust artefact.) Lateral projections of stress maps (bottom) showing individual protopodia interdigitated by naked cuticle. As a contractile wave (grey box) progressed through the animal, protopodia were lifted off the substrate. Scale bar=100µm. **b**, WARP image sequence of larva prior to (–1.5s to –0.5s) and engaging in (0s) a headsweep (representative of 2 animals and 3 turns). Note the large posterior displacement (blue arrow)(Images cropped around the animal). Scale bar=200µm. **c**, Profiles of cavity displacement along anteroposterior (A-P) axis in resting state (black dotted line at –1.5s in **b**) and pre-headsweep (red dotted line at –0.5s in **b**), showing that peak displacement decreased across all segments from the resting state (grey box) to pre-headsweep (pink box). **d**, Bilateral displacement profile across the mediolateral (ML) axis of the A4 protopodium (solid lines in **b**) at different times prior to the headsweep, showing that the width of the contact increases from the resting state (– 1.5s) to the pre-headsweep state (–0.5s) and partially reduces again immediately after head movement. **e**, (**i**) Brightfield image (3^rd^ instar larva) and (**ii**) displacement map (2^nd^ instar larva) of the posterior-most body segment, showing how two cuticular protrusions (white arrowheads) and the terminal protopodium (A8) generate a tripod-shaped substrate displacement. (**iii**) Profiles along blue and red dotted lines in (ii). Scale bar=200µm (i) and 100µm (ii). **f**, Sequence of displacement maps of tripod structure before the start of a forward wave (<0.24s) and the removal of tripods upon beginning of peristalsis (>0.48s). Scale bar=100µm. **g**, Percentage of forward waves (FW), bilateralisms (BL), backward waves (BW) preceded by tripod contact, and tripod deployments without any observed locomotor behaviour (unrelated). **h**, Time delay between tripod deployment and initiation of movement at A7. Points colour-coded by animal, n=6. Line=mean, box=±1 standard error of the mean, whiskers=±1 standard deviation.

We also used WARP to investigate the bilaterally asymmetric head sweeps generated by *Drosophila* larvae to sample odours and direct navigation. During head sweeps, anterior segments and mouthhooks detached or dragged across the substrate before replanting (**Figure 4B**). 0.5-1s prior to headsweep initiation, the contact area in posterior segments increased, spreading outwards laterally, employing both the protopodia and the naked cuticle along the midline (**Figure 4C**). This broad but shallow anchoring quickly returned to the ordinary resting phase profile after the mouth hooks were replanted onto the substrate (**Figure 4D**).

Before forwards waves and headsweeps, larvae produced large indentations posterior to their terminal segment. Anatomical examination revealed accessory structures located at the terminus of the posterior abdomen. Together with the terminal denticle band, these cuticular processes generated tripod-shaped indentation patterns (**Figure 4E**). The left and right sides of the tripod deployed and detached simultaneously (**Figure 4F**). Tripod formation was seen before all observed forwards waves (n=28 across 6 animals) and bilateral thoracic activity (n=3 across 2 animals), but not all tripod contacts resulted in further behaviour (**Figure 4G**). To investigate the relationship between tripod placement and locomotion further, we recorded the delay between tripod contact and protopodial detachment in A7. The mean delay was 0.66s*±*0.21s (**Figure 4H**, n=20 waves across 6 animals).

Next, to estimate the GRF associated with the indentation of each protopodium, we integrated the displacement and stress maps over the region covered by each protopodium. During forward waves, the temporal evolution of GRFs mirrored the characteristics of the cycle seen in the stress maps, with absolute GRFs ranging between 1 and 7µN (**Figure 5A**). However, unexpectedly, we observed an additional force applied to the substrate both when protopodia leave the substrate (SI) and when they are replanted (ST). To investigate whether this force was due to an active behaviour or due to shifting body mass, we plotted protopodial GRFs against the contact area for each protopodium over time, combining data from multiple forwards waves (**Figure 5B**). We found that the magnitude of force output was positively correlated with protopodial contact area in a quadratic relationship (A6: Adj. R^2^=0.77, A4: Adj. R^2^=0.92, A2: Adj. R^2^=0.79) Comparing different animals, we find that GRFs were relatively consistent across most segments (**Figure 5C**).

**Figure 5.**
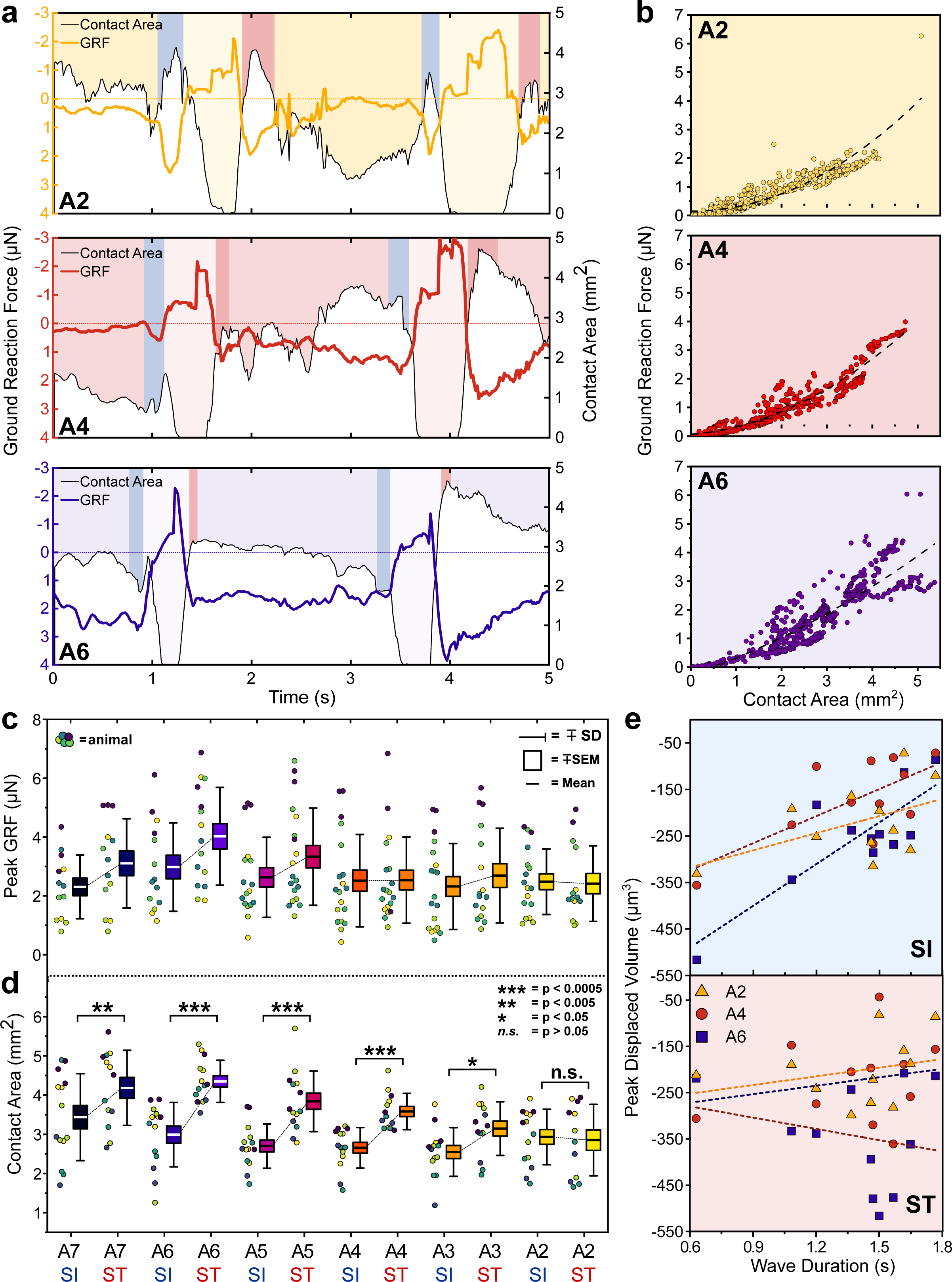
Protopodia produce GRFs in the micronewton range and show complex spatiotemporal dynamics. **a**, Ground reaction force (GRF, coloured line) and protopodial contact areas (white area under black line) during forward crawling for A2, A4 and A6 protopodia, showing progression of waves through animal (light-coloured boxes). Blue (SI) and pink (ST) boxes denote characteristic troughs in GRF immediately prior to protopodia leaving the substrate and returning to the substrate, respectively. **b**, GRFs exerted by different protopodia show a 2^nd^ order polynomial relationship (dashed line) with the contact area of that protopodium (A6: Adj. R^2^=0.77, A4: Adj. R^2^=0.92, A2: Adj. R^2^=0.79). Peak GRFs and **d**, peak contact area during SI and ST across body segments. Data points denote single events, colours indicate different animals. n=5, 15 waves. Contact areas were compared by a two-way repeated measures ANOVA (*<0.05, **<0.005, ***<0.0005, n.s.=not significant). **e**, During SI, peak displaced volume scaled with wave duration for larger abdominal segments (A6: R^2^=0.69; A4: R^2^=0.48) but not for smaller anterior segments (A2: R^2^=0.24). During ST, displaced volume did not scale with wave duration regardless of the segment (A6: R^2^=0.05; A4: R^2^=0.05; A2: R^2^=0.08). n=4, 11 waves.

The contact area of each protopodium showed a pronounced peak during SI and ST. The maximum contact area during ST was significantly greater than during SI for the posterior abdomen (p*≤*0.05 for A8/9-A3) but not for the anterior abdominal protopodium (p>0.05 for A2) (**Figure 5D**). The peak of the displaced volumes during SI was largely determined by wave duration (R^2^ range: 0.48-0.69, A7-A4, **Figure 5E**), again except for the anterior abdomen (A3: R^2^=0.15; A2: R^2^=0.24). However, the peak of the displaced volumes during ST did not scale with wave duration (R^2^ range: 0.03-0.05, A7-A2). This suggests that protopodia push off from the substrate harder during faster waves, but that varying wave speed does not strongly influence forces exterted onto the substrate during protopodia placement. This observation is consistent with our morphometric data, which showed that wave duration is associated with SI latencies but not with ST latencies.

### Sub-protopodial force dynamics

Lastly, to investigate how forces are translated into the substrate within a single protopodium during a ‘footfall’ cycle, we examined the spatiotemporal substrate interaction during the ST (**Figure 6A**). This showed how protopodia expand their indentive contact across both the AP and mediolateral (ML) axes when being replanted. Kymographs along the AP midline of animals and profiles running up the AP axis extracted from these revealed a delay between when the most posterior and the most anterior part of the protopodium contacts the substrate (**Figure 6B**). The mean contact delay relative to the most posterior part of the protopodium was 0.035s*±*0.007s at 6µm away from the most posterior part and increased to 0.062s*±*0.021s and 0.253*±*0.115s in the middle and at the most anterior part of the protopodium, respectively (**Figure 6C**).

**Figure 6.**
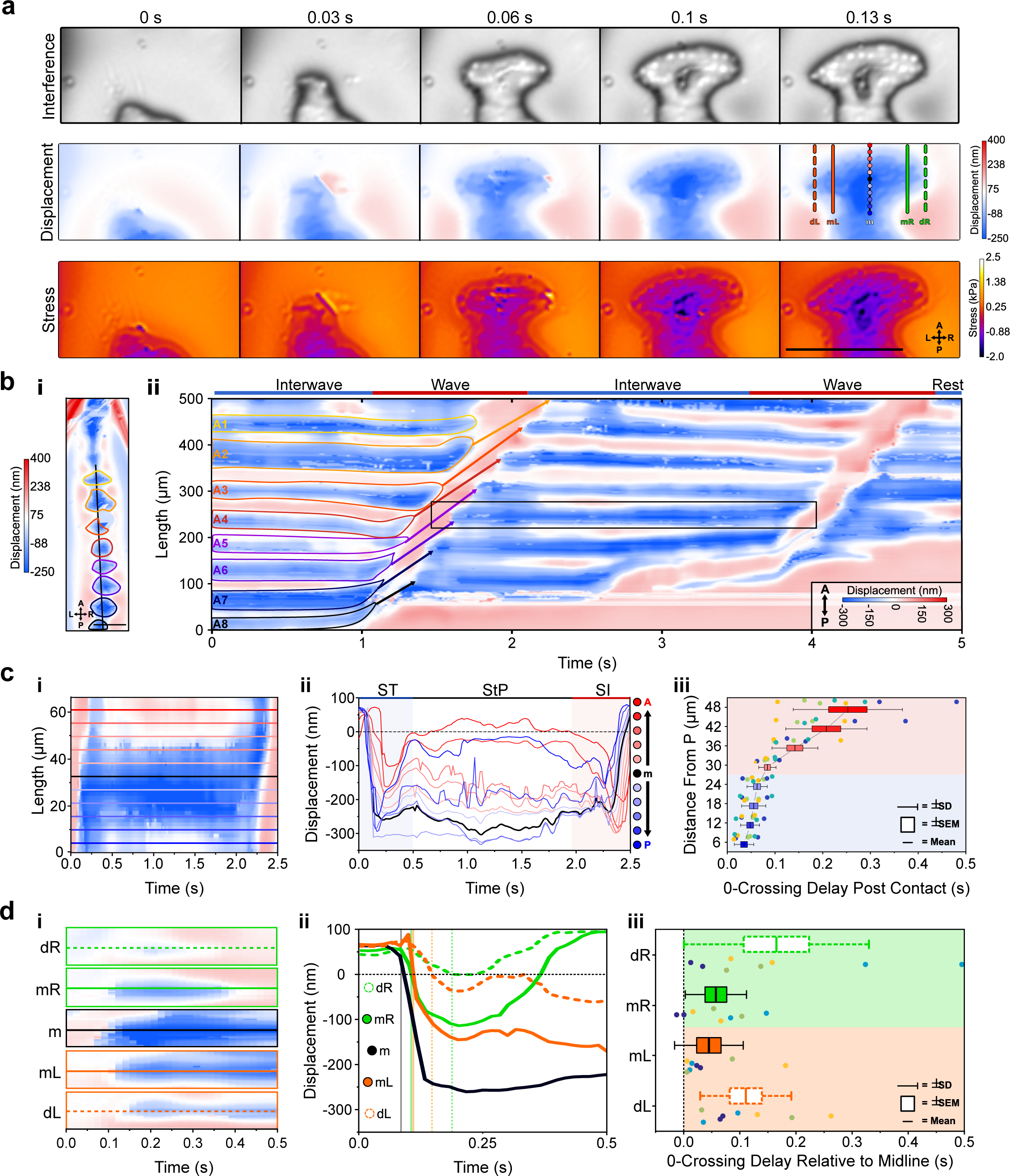
Sub-protopodial force dynamics reveal sub-step processes and functional substrate interfacing domains in each step. **a**, WARP imaging of protopodial landing during ST of an A6 protopodium. Raw interference images from WARP acquisition show footprints of individual denticles as white dots. Displacement and stress maps show how landing starts with posterior denticle rows before spreading out along the AP and ML axes. Scale bar=100µm. **b**, **(i)** Displacement map of whole animal. **(ii)** Kymograph of displacement along AP axis (black line in i) over 2 forward waves. Bands of red and blue correspond to naked cuticle and protopodia, respectively. Scale bar=100µm. **c**, (**i**) Kymograph of displacement along the AP axis of an A6 protopodium (box in **b**). (**ii**) Profiles across kymograph at different positions along the AP axis of protopodium (lines in i). (**iii**) Latency of substrate indentation (displacement <0nm) during ST along the AP axis, relative to the extreme posterior of protopodium. Compared to the posterior half of protopodium (light blue area), the anterior half shows larger latencies and variations in latency (light red area). n=4 animals, 8 swing termination events. **d**, Kymograph of displacement along AP axis during ST for the distal left (dL), medial left (mL), midline (m), medial right (mR) and distal right (dR) section of the A6 protopodium. Height of each kymograph, 66.42µm, (**ii**) Profiles across the central AP line of each kymograph in (i). Vertical lines indicate times when midline, medial right/left and distal right/left indentation starts (displacement <0nm). (**iii**) Latency of substrate indentation during ST relative to the midline for medial right/left and distal right/left locations. n=4, 8 swing termination events.

To examine how protopodia expand along the ML axis, we performed a similar analysis, taking kymographs and profiles for the displacement maps at different distances to the midline of a protopodium. At a medial distance from the midline, the contact delay relative to the midline was 0.045s*±*0.022s (left) and 0.057s*±*0.019s (right). At the distal left and right of the protopodium, contact occurred 0.111s*±*0.030s (left) and 0.165s*±*0.058s (right) after midline contact (**Figure 6D**). This analysis also showed that protopodia insert a medial-spike into the substrate, through which ST related peak GRFs are conferred, before expanding along the AP and ML axes.

## Discussion

### *Drosophila* larvae, though legless, have protopodia

The cuticle of larvae shows distinct patterns of denticulation (denticle bands) and the developmental processes which give rise to these features have been well studied (33) though their role in locomotion has long been unclear (47). Here, we show that denticle bands are situated upon larger articulated foot-like cuticular processes, which act as locomotory appendages. Protopodia dynamically change shape during locomotion, allowing sequestration and presentation of denticles. Individual protopodia and individual denticles exert GRFs in the 1-7µN and 1-48nN ranges, respectively. Superficially, protopodia resemble the much smaller pseudopodia in cells – transient structures, similarly covered with actin protrusions, used by cells to facilitate movement (48). The same function and principles of protopodia may underlie ‘creeping welts’ noted in larger dipteran larvae (49) and show similarities to soft prolegs of *Manduca sexta* caterpillars but are approximately 30 times smaller (8).

### Insights from morphometric kinematic tracking of denticle band movements

Our study provides, to our knowledge, the first detailed description of the morphometry of denticle bands during movement, showing how denticle bands are deployed onto and removed from the substrate. Posterior denticle rows hit the substrate before anterior rows during deployment (ST) and left the substrate before anterior rows during removal (SI). This suggests that both deployment and removal involved rolling ‘heal-toe’ like movements, similar to footfalls in limbed animals, including terrestrial arthropods (40). Removal but not deployment correlated with wave duration. In *Manduca* caterpillars, it has been noted that SwPs scale positively with wave duration (50); however, to our knowledge there is no measurement for SI and ST in these animals.

SI latencies scaled positively with wave duration across most segments whereas ST latencies did not show this trend. SIs scale with SwP and this could be mediated by proprioceptor activity in the periphery (51). Fine sensorimotor control of musculature during this process would allow for precisely tuned propulsion during peristalsis. In contrast, the more random nature of the ST suggests the process is less finely controlled. This could be a consequence of fluid inertia within the animal, and/or the release of elastic energy from cuticle (38) or relaxation of muscles (50,52).

### ERISM-WARP allows computation of ground reaction forces in *Drosophila* larvae

We adapted state-of-the-art mechanobiological force measuring techniques to enable measurement of substrate interaction dynamics of a freely behaving soft-bodied animal with micrometer spatial resolution, millisecond temporal resolution and nanonewton force resolution. Previously, high-resolution force mapping was limited to cellular mechanobiology. Specifically, we developed microcavity resonators tuned to the vertical forces generated by larvae and employed ERISM and WARP to perform direct measurements of substrate interactions in anesthetized and behaving animals. GRFs produced by individual denticles in anaesthetised animals were in the ∼11nN range. The measured vertical GRFs produced by the individual protopodia of each segment were in the 1 to 7µN range, roughly three orders of magnitude less than the 17mN recorded from an entire 1.72g *Manduca sexta* caterpillar (8). Our measurements provide fundamental constraints for future biomechanical modelling studies seeking to incorporating these structures.

Displacement and stress maps produced during larval crawling revealed that animals can control when and how protopodia contact the substrate. We observed that larvae travel surrounded by moisture from a water droplet, which produces a relatively large upwardly directed force in a ring around the animal. This surface tension produced by such a water droplet likely serves a role in adhering the animal to the substrate. However, during forward waves, we found that protopodia detached completely during SwP, suggesting this surface tension-related adhesion force can be easily overcome by the behaving animal. This observation, coupled with our lateral imaging of protopodia in constrained animals, explains how larvae prevent their rough denticulated cuticle from creating drag due to friction against the direction of the wave. Larvae do not simply pull protopodia off the substrate in a vertical direction; instead, they horizontally slide posterior regions forward in the axis of travel, before invaginating and therefore sequestering friction generating features (e.g. denticles). This shows similarities to the use of shearing forces to detach adhesive pads in limbed arthropods (40) Inversion of the cuticle to remove denticles from the substrate may also explain why natural variations in denticle count across animals do not strongly affect locomotor behaviour (47). The invagination process is reversed in order to expand the protopodia into and locally across the substrate, providing an expanding anchor which can serve as a postural support to enable locomotion and prevent lateral rolling during bilaterally asymmetric behaviours such as head sweeps. The dynamic anchoring during the progression of peristaltic waves thus serves to counteract horizontal reaction forces resulting from Newton’s 3^rd^ law of motion. Such a sequence of positioning points of support and anchoring them against the substrate has long been postulated to be a fundamental process in soft-bodied locomotor systems (6); and may be central to explaining why soft-bodied animals have evolved segmentally repeating bodies (53). However, WARP microscopy is largely limited to measurements of forces in the vertical direction, and though we can make inferences such as this as they are a consequence of fundamental laws of physics, we present this conclusion as a testable prediction which could be confirmed using a force measurement technique more tuned to horizontally directed forces relative to the substrate.

Our ERISM-WARP measurements also revealed substrate interaction from accessory structures. Immediately before enacting headsweep, larvae redistributed their body mass into naked cuticle in between protopodia along the midline, effectively fusing multiple protopodia into a single ‘ultra-protopodia’ that extends across multiple posterior segments. This redistribution occurs hundreds of milliseconds before the start of a head sweep, suggesting that it may be part of an active preparatory behaviour. Similar preparatory behaviours have been observed in caterpillars before cantilevering behaviours (10), adult flies during fast escape behaviours (54) and humans during stepping (55). More detailed characterisation of this behaviour remains a challenge owing to the changing position of the mouth hooks. Due to their rigid structure and the relatively large forces produced in planting, mouth hooks produce substrate interaction patterns which our technique struggles to map accurately due to overlapping interference fringes ambiguating the fringe transitions.

We also observed transient tripod-shaped substrate interactions in posterior terminal regions of larvae immediately before forward waves and headsweeps. Two bilateral cuticular protrusions covered in trichomes, labelled in previous work as anal papillae (56), are likely candidates responsible for these substrate interactions. However, the actions of these structures have hitherto not previously been described as a part of movement in soft-bodied animals. Each body segment has a preceding substrate-planted segment which acts as the anchor and lever to push the animal forward. However, A8 is an exception; it has no full preceding segment in contact with the substrate to counteract its muscle contraction. The tripod processes are ideally positioned to provide an anchor against horizontal reaction force generated by by the initial contraction when moving forward (**Figure 7A**), and might effectively form a temporary extra segment prior to initiation of a wave (**Figure 7B**). The deployment of cuticular features as transient anchors has not been a focus of previous studies; future work should incorporate our findings into models of crawling behaviour. WARP and ERISM have technical limitations, such as the difficulty of resonator fabrication. This problem is compounded by the fragility of the devices owing to the fragility of the thin gold top mirror. This becomes problematic when placing animals onto the microcavities, as often the area local to the initial placement of the animal is damaged by the paintbrush used to move the animals. Further, as a result of the combining of the two wavelengths, the effective framerate of the resultant displacement and stress maps is equal to half of the recorded framerate of the interference maps. This necessitates recording at very high framerates and thus requires imaging at reduced image size to maximise framerates, but this in turn reduces the number of peristaltic waves recorded before the animal escapes the field of view. A further limitation is that WARP and ERISM are sensitive mainly to forces in the vertical direction; this is complementary to TFM, which is sensitive to forces in horizontal directions. Using WARP in conjunction with high speed TFM (possibly using tuneable elastomers presented here) could provide a fully integrated picture of underlying vertical and horizontal traction forces during larval locomotion.

**Figure 7.**
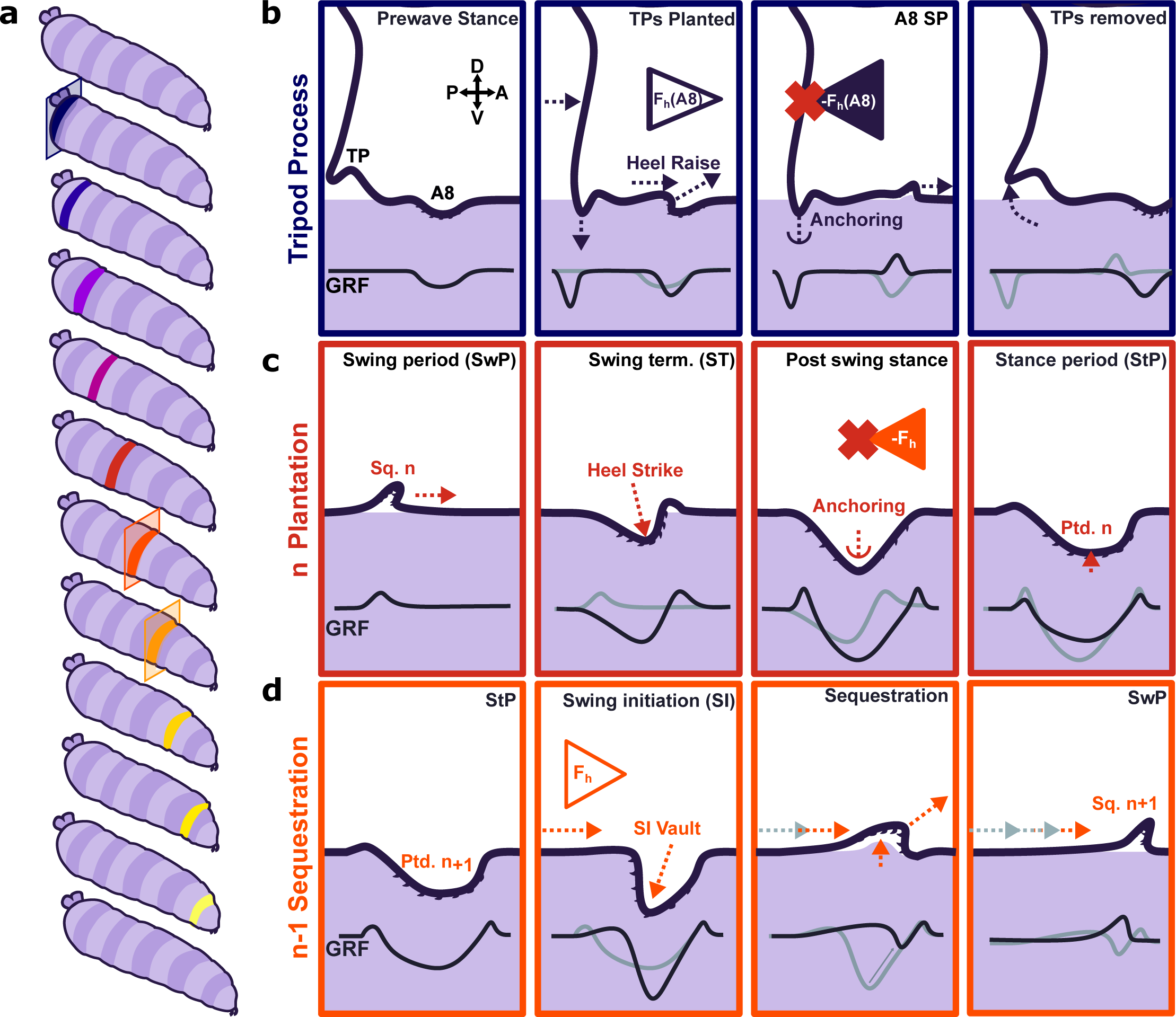
Proposed model for protopodia-substrate interactions during *Drosophila* larval locomotion. **a**, Schematic illustration of forward wave propagating from posterior (blue) to anterior (yellow). **b**, At the start of a forward wave, animals contract the posterior-most abdominal segment (A8), producing an anterograde horizontal force *F*_h_(A8). Due to Newton’s 3^rd^ law, there is an equal but opposite reaction force –*F*_h_(A8). To counteract this force, tripod processes (TPs) deploy onto the substrate and generate a temporary anchor, allowing the A8 protopodium to swing forward. **c**, During swing termination (ST) at the end of the swing period (SwP) of segment *n*, the corresponding sequestered protopodium (Sq. *n*) strikes the substrate with its posterior most denticle row, then gradually unfolds into the substrate along its entire anteroposterior extent. During the stance period (StP), this planted segment *n* (Ptd. *n*) forms an anchor to mitigate the retrograde reaction force due to the subsequent contraction of segment *n*-1. **d**, In time with anchoring of protopodium *n*, protopodium *n*-1 performs swing initiation (SI) by removing denticles from the substrate and sequestering into an invagination pocket, which reduces friction during the subsequent SwP. The contraction of segment *n*-1 then leads to an anterograde force (*F*_h_) that is balanced by the anchoring of protopodium *n* as illustrated in **c**.

### Evidence for functional subdivisions within protopodia

By examining the dynamics of individual footfalls, we found that protopodia exhibited characteristic spatiotemporal force patterns across the footfall cycle. This shows parallels to the regional specificity of function in a vertebrate foot. Specifically, the posterior medial region of the protopodia makes a large contribution to peak GRFs exerted during ST (**Figure 7B**), similar in nature to a vertebrate heel strike impacting the surface prior to the rest of the foot. We propose that this zone of the protopodia acts as a vaulting point for the protopodia, functioning as a ‘point d’appui’ (point of support) as proposed in other soft-bodied animals (6,57). The transience of this vaulting point suggests it may be critical for locomotion, but dispensable for postural control during StP. The distal area of protopodia exhibited a similar transience. This increased force transmitted into the substrate is unexpected as the forces generated for the initiation of movement should arise from the contraction of the somatic muscles. We propose that the contraction of the musculature responsible for sequestration acts to move haemolymph into the protopodia thus exerting an increased pressure onto the substrate while the contact area decreases as a consequence of the initiation of sequestration. Immediately after the posterior and medial protopodia impact during ST, the contact area of the outer region of the protopodia grew across both the AP and the ML axes. However, throughout the StP, this outer region then slowly retracted, suggesting it too was not critically important for maintaining posture during StP. This may reflect a transient anchoring mechanism – specifically, this anchor region deploys to provide greater friction for the subsequent segments (**Figure 7C**). This would allow the contractile wave to progress unimpeded by resultant reaction forces. Previously, such a function was thought to be provided mainly by mucoid adhesion (6). However, *Drosophila* larvae are proficient at crawling over wet surfaces where mucoid adhesion is reduced or impossible (43). Larvae can adhere to dry surfaces but have difficulty moving over these, although mucoid adhesion would provide optimal anchorage in this context. Water surface films appear to facilitate larval locomotion in general, but the biomechanical mechanisms by which this occurs remain unclear. We propose that protopodia act to provide an optimal balance between anchorage and adhesion depending on the environmental context. Overall, our work suggests that *Drosophila* larvae use a sophisticated process of articulating, positioning and sequestering protopodia to enable movement over terrain. Future work will be needed to determine the extent to which these processes are conserved across other soft-bodied crawlers.

### Conclusions and outlook for future work

Combining ERISM-WARP with a genetically tractable model organism opens new avenues for understanding the biomechanical basis of animal behaviour, as well as the operation of miniaturized machines. Here we have provided new insights into the relatively well-studied behaviour of *Drosophila* larval locomotion. We have provided new quantitative details regarding the GRFs produced by locomoting larvae with high spatiotemporal resolution. This mapping allowed the first detailed observations of how these animals mitigate friction at the substrate interface and thus provide new insights into how locomotion is achieved in soft animals. Further, we have ascribed new locomotor function to appendages not previously implicated in locomotion in the form of tripod papillae, providing a new working hypothesis for how these animals initiate movement. It is our hope that these new principles underlying locomotion outlined here serve as useful biomechanical constraints as called for by the wider modelling community (39) We used *Drosophila* larvae as a test case, but our methods now allow elastic optical resonators to be tuned to a wide range of animal sizes and thus create new possibilities for studying principles of neuro-biomechanics across an array of animals. In parallel, roboticists are increasingly moving to create miniaturized soft robots for a variety of applications. Our approach is well suited to provide ground truth, constraints, and inspiration for the development of such miniaturized machines. It also provides a potentially powerful new resource for evaluating the performance of these devices, as our methodology will also allow scientists to measure GRFs during the operation of miniaturized soft machines. Importantly, while we have focused here on the movement of soft animals, our sensors could also be tuned to measure forces produced by small limbed animals or miniaturized machines with rigid internal or external skeletons. Overall, this work therefore establishes a flexible platform for future investigations aimed at integrating knowledge across genetics, neuroethology, biomechanics, and robotics.

## Materials and Methods

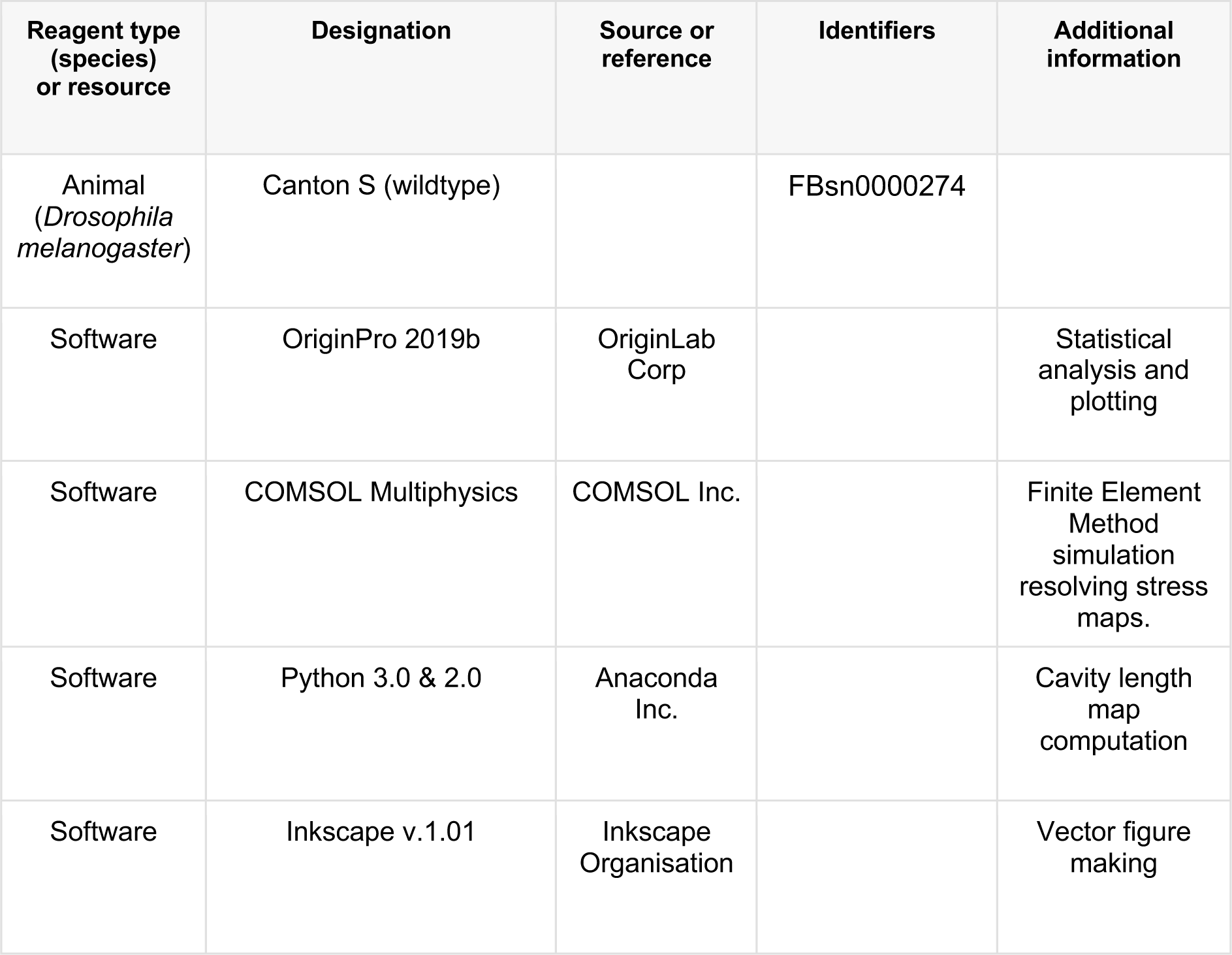
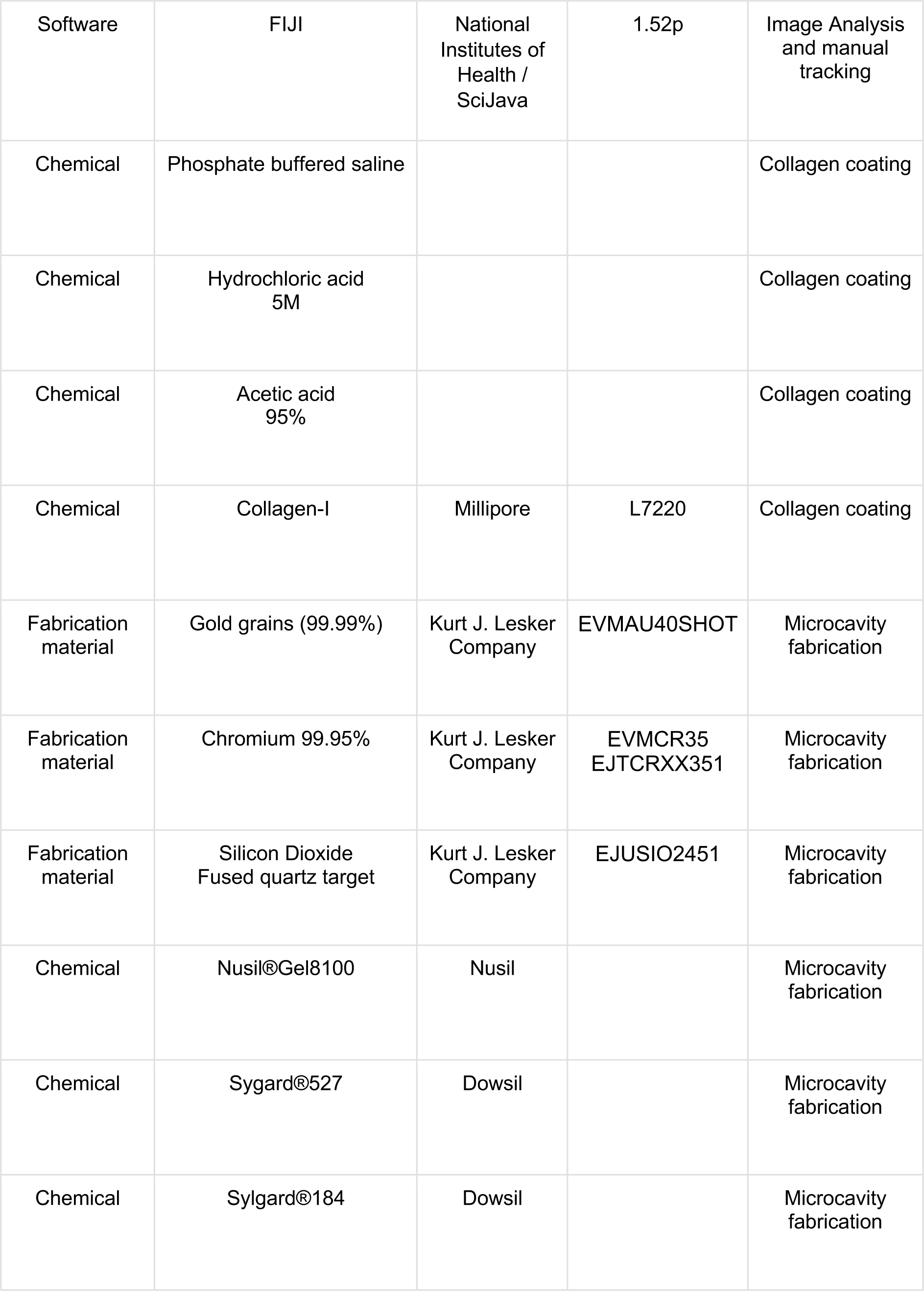
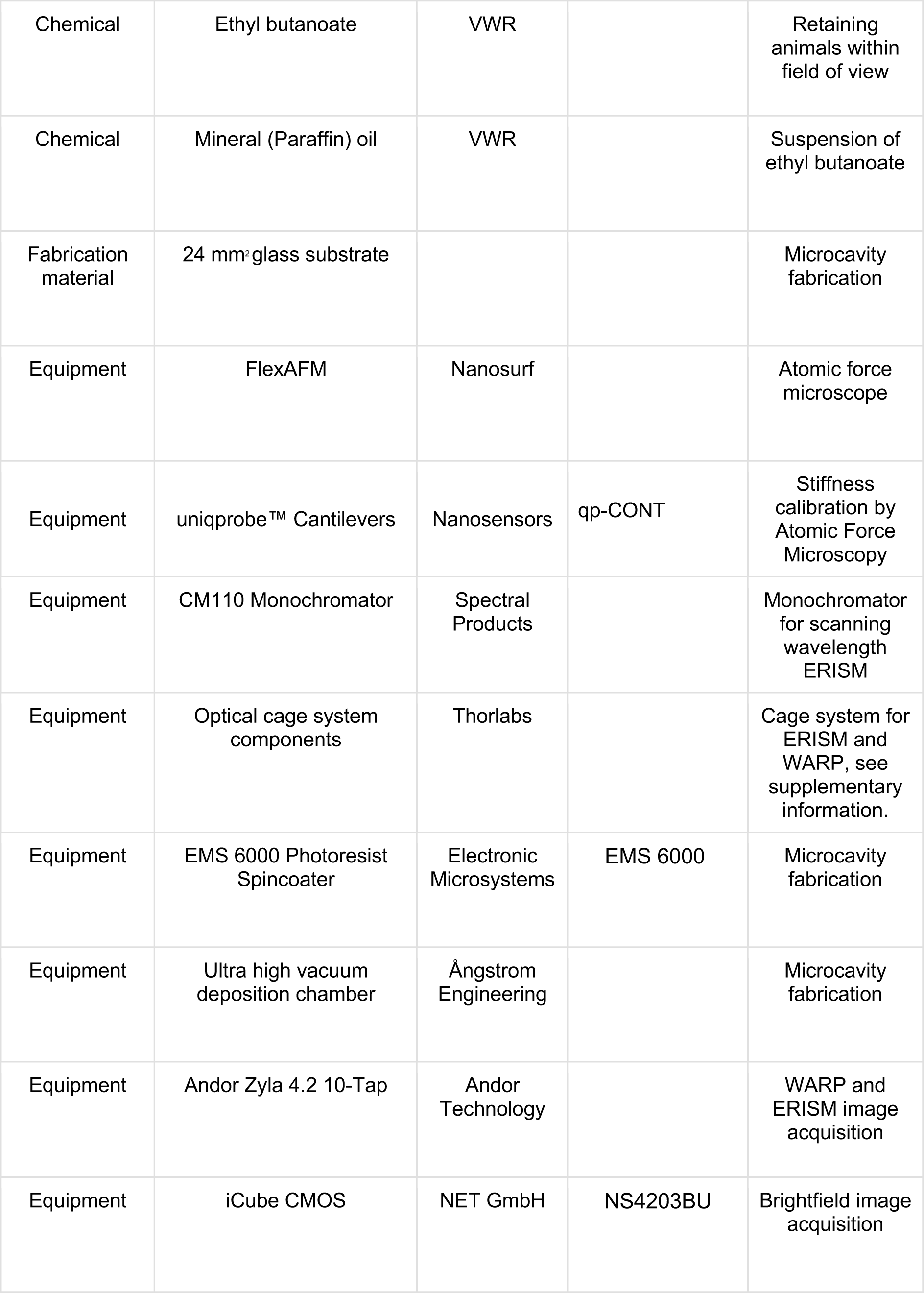
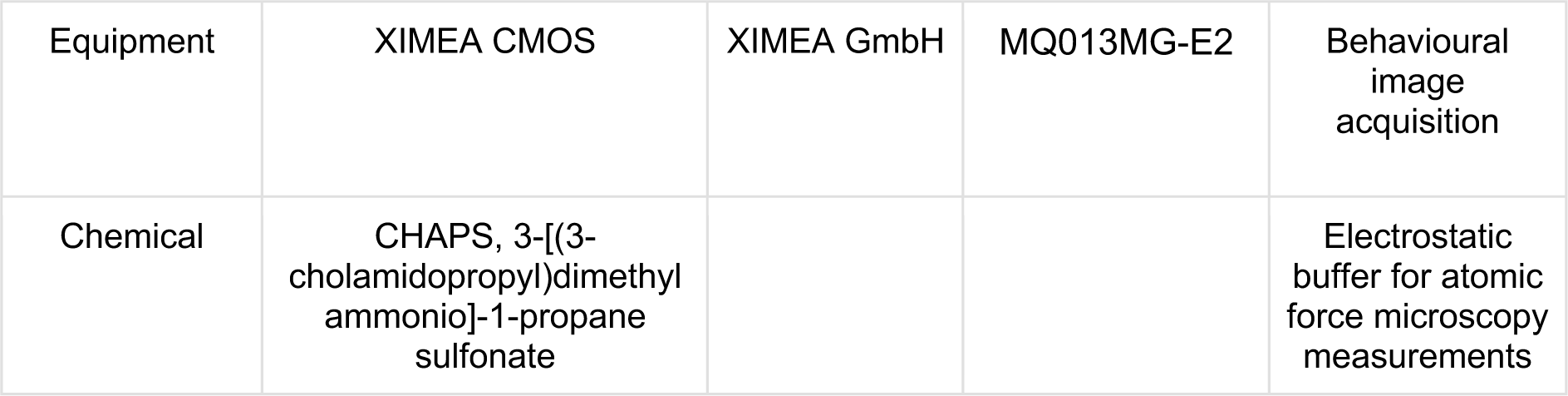
Key Resources.

## Animal Rearing

Animals were raised on standard cornmeal and yeast medium (17.4 g/L yeast, 73.1 g/L cornmeal, 5.8 g/L agar, 4.8 ml/L propionate) at 25°C with a 12-hour light-dark cycle except where explicitly stated otherwise. Animals were given at least 1 hour to acclimate to room temperature prior to all experiments. Immediately prior to experiments, samples of media containing larvae were taken using a spatula before being placed into a columnar stacked sieve with 40, 60 and 100 meshes from top to bottom, respectively. Media samples were run under gentle flowing tap water to separate adult debris, 2^nd^ instar larvae and 1^st^ instar larvae with embryos on each mesh. Larvae from the 60-mesh fraction of the sieve were observed under a microscope and animals around 1 mm were selected and washed before being placed on 1% (w/v) agarose lined dishes.

## Microcavity fabrication

The fabrication protocol of elastic microcavities was adapted from (58). 24 mm^2^ borosilicate glass substrates of No.5 thickness were cleaned via ultrasonication in acetone followed by propan-2-ol for 3 minutes. After cleaning, substrates were dried using N_2_ and baked at 125°C for 10 minutes to clear any residual solvent. Cleaned glass substrates were then plasma treated with oxygen plasma for 3 minutes at 20 SCCM O_2_ flow rate to clear any residual organics and activate the surface of the glass. Cleaned and activated glass substrates were then sputter coated with 0.5nm of Cr, which acted as an adhesion layer for the subsequent 10nm Au layer which was deposited by thermal vapour deposition. 50nm of SiO_2_ was then deposited by sputter coating to improve stability of the resultant bottom mirrors. Roughly 100µl of pre-mixed and degassed polydimethylsiloxane gels was spincoated onto the bottom mirrors at 3000RPM, 1500RPM acceleration, for 60 seconds and then quickly transferred to a pre-heated metal plate at 150°C for 1.5 hr to cure the elastomer. After curing, elastomer coated bottom mirrors were O_2_ plasma treated with the desired plasma power at 20 SCCM O_2_ flow rate for 10 seconds. 15nm of Au was then deposited onto the oxidised elastomer, thus completing the microcavity.

## Microcavity characterisation

Microcavities were characterised using a NanoSurf Flex Atomic Force Microscope (Nanosurf, Liestal, Switzerland). 15-18µm diameter glass beads were glued to the tip of Uniqprobe™ QPCont cantilevers (Nanosensors AG, Neuchatel, Switzerland) using a UV-polymer glue after thermal calibration of the spring constant at 21°C. Sphere-tipped cantilevers were then indented into microcavity samples at 1µm/s with up to 30nN of force. This process was repeated across the surface of the microcavity at least 5 times, with each measurement being roughly 2mm apart to get a measure of the variation across the cavity surface. Force-distance profiles recorded by the AFM were then fitted to the Hertz model to compute the Young’s modulus at each point of each sample. Mean cavity lengths were measured by taking 4 ERISM images at 4x magnification from each corner of the cavity, and then taking the mean of 4 regions of interest per image.

Prior to use in experiments, a 12-well silicone chamber (ibidi GmbH, Munich, Germany) was cut such that only 1 large square-well, originally comprised of four smaller wells cut off from the rest of the chamber, remained and was placed onto a microcavity. A low pH Collagen-I (1mg/ml; Millipore L7220) solution was then prepared at a 1:1 (v/v) ratio with pH3 phosphate buffered saline (PBS). pH3 PBS was prepared with either hydrochloric acid or acetic acid, mixing until pH3 was recorded using an electronic pH meter. Collagen-I mixtures were then dosed onto microcavities in silicone wells (1ml per microcavity) and allowed to coat the surface overnight at 4°C. Immediately before the experiment, microcavities were washed with deionised water at least 5 times, taking care not to remove all liquid to prevent damage to the top gold surface.

## Denticle band kinematic imaging

All animals were raised in ambient light conditions at room temperature. Between 48 and 72 hours after flies were introduced to fresh media, feeding 2^nd^ instar Canton-S wildtype animals were selected with a size exclusion criterion – any animals below 0.8 mm or above 1.5 mm were rejected. Animals were then washed and allowed to acclimate to 0.5% (w/v) agarose.

Immediately before experiments, a single animal was transferred to a freshly set dish containing 0.5% (w/v) agarose while still transparent. These dishes were then quickly placed onto the 3D printed stage of a custom-built inverted Bresser Advance ICD stereomicroscope (Bresser GmbH, Rhede, Germany). Denticle band images were acquired, through the still transparent agarose substrate, at 60 frames per second for at least 1 minute while the larva was freely behaving. All images were acquired using a XIMEA CMOS camera (XIMEA GmBH, Münster, Germany) through MicroManager 1.4 (59). The velocity of 33 individual identifiable points across the animal’s body during peristaltic waves whilst imaging from the ventral side of 2^nd^ instar larvae Denticle bands were tracked manually using the Manual Tracking plugin of ImageJ (60). Analysis of tracking data was performed using OriginPro 2019 (OriginLab Corporation, MA, USA).

## ERISM and WARP imaging

Elastic Resonator Interference Stress microscopy (ERISM) was used to record high-resolution maps of substrate indentations by monitoring local changes in the resonances of a soft and deformable optical microcavity. ERISM has been used to quantify cellular forces down to the piconewton range. The static thickness of microcavities was measured adapting our previously published ERISM method as described in (24,58). In brief, images of the cavity were taken under epi-illumination with a series of 201 different wavelengths (550-750nm in 1nm steps). From these images, the minima in the spectral reflection for each pixel were correlated with theoretical values obtained from optical modelling for cavities of different thicknesses to determine the actual thickness at each position across the image (Cavities were between 8-12µm in static thickness.) Thickness maps were converted into maps of local displacement by subtracting a linear plane, using the mean thickness of the cavity in each corner.

For dynamic force mapping, we used a further improved version of the WARP routine described in (26). Epi-illumination with light of two different and quickly alternating wavelengths was produced by passing the emission from two identical red LEDs (dominant emission wavelength 625nm, FWHM 17nm; Thorlabs Inc. NJ, USA) through two identical narrow bandpass filters (peak transmission at 633nm, FWHM of 1nm; Thorlabs Inc., NJ, USA). By tilting the filter located in front of one of the LEDs by approximately 15 degrees relative to the incident light, its peak transmission wavelength was tuned to λ_θ_=628nm. For the optical modes supported by our microcavities, this corresponds to a phase shift of roughly 90°, but remains within the same free spectral range band of the cavity. For the WARP measurements, we first took calibration images (under subsequent illumination at λ and λ_θ_) of the empty microcavity in an area with roughly linear slope in cavity thickness, e.g. near where the silicone well containing the larvae meets the surface of the cavity. Images of behaving larvae were then recorded under rapidly alternating illumination at λ and λ_θ_, with the camera sending alternating trigger pulses to each LED to generate interleaved stacks of λ and λ_θ_ images.

Displacement maps were obtained from these stacks using a series of image transformations, based around the fact that the ratio of the difference and the sum of pixel intensities at λ and λ_θ_ is linked to local thickness in an unambiguous manner, at least across each free spectral range. See Supporting Information Fig. S4 and (26) for further details on the calculation of displacement from the λ and λ_θ_ images. All WARP and ERISM images were acquired using an Andor Zyla 4.2 sCMOS camera (Andor Technology, Belfast, UK).

Stress maps were calculated from the ERISM and WARP displacement maps as described previously (23), using a finite element method simulation via COMSOL multiphysics (COMSOL Ltd., Cambridge, UK) and the known mechanical properties of the microcavity.

## Polydimethylsiloxane gel preparation

Polydimethylsiloxane elastomers were prepared according to manufacturer guidelines for all gels. The two component precursors of different gels were mixed together in separate glass bottles, using an equal mass ratio of the two components for Sylgard 527 and NulSil Gel8100 but a 1:10 volumetric ratio for Sylgard 184. Mixing was performed by 10 min of magnetic stirring (Sylgard 527 and NuSil GEL8100) or by 10 min of mechanical stirring (Sylgard 184). The elastomer mixtures were then combined in a fresh bottle in the desired mass ratio using a syringe following the same method as a previous study (44). Combined elastomers were mixed for a further 10 minutes. Mixtures containing Sylgard®184 were initially mixed by high speed vortexing to coarsely disperse the gel to allow for the magnetic stir bar to overcome the high viscosity of the gel. After mixing, all preparations were degassed under vacuum for around 5 minutes, prior to fabrication of microcavities.

## Anaesthetised animal force imaging

Animals were selected, cleaned, and placed in a fridge at 4°C for 2-3 hours to anaesthetise them. Immediately prior to experiments, anaesthetised animals were gently placed onto a collagen coated microcavity in a petri-dish on ice. The microcavities were then placed, using a moistened paint brush, on the ERISM-WARP microscope and the animals were observed carefully. As soon as mouth-hook movement was observed, an ERISM measurement was taken. Animals often had to be placed back onto ice to anaesthetise them once more as they rapidly regained motility. As the complete ERISM scan requires ca. 5 seconds, animals were required to be completely stationary in order to obtain reliable stress map images.

## Freely behaving animals force imaging

Animals were selected according to the previously outlined criteria and cleaned before being placed onto a 1% (w/v) agarose lined petri dish. Elastic resonators were prepared according to the coating criteria mentioned above. 10% Nusil®GEL8100, 180W O_2_ plasma treated microcavities were used for all freely behaving experiments. Once calibration images of the microcavity were acquired, excess water was removed from the cavity and animals were gently placed onto the cavity surface with a paintbrush, taking care to ensure there was enough moisture on the animal to prevent drying by wetting the paintbrush prior to transferring the animal. In order to keep animals on the sensor surface, a 50µl drop of 15mM ethyl butanoate (Sigma-Aldrich Inc., MO, USA), suspended in paraffin oil, was dropped onto a 24 mm^2^ glass coverslip before being inverted and placed on top of the silicone well (ibidi GmbH, Munich, Germany) such that the attractive odorant faced towards the animal but perpetually out of its reach. Animal substrate interaction was then imaged by WARP, using alternating wavelengths to generate a series of interleaved cavity resonance images, and displacement and stress maps were generated as described prior. All WARP videos were recorded at 120FPS, producing displacement maps with an effective framerate of 60FPS, using a 4X magnification objective. Due to the high framerate, we were limited to the use of ¼ of the total camera sensor, thus higher magnifications would prevent mapping of the whole-animal.

## Statistical Analyses

All statistical analysis was performed using OriginPro 2019 (OriginLab Corporation, MA, USA). Coefficients of determination (R^2^) for all but GRF vs contact area analysis were determined using a linear fit. The rarity of backwards waves during normal larval behaviour precluded analysis of latencies as used in **Figure 2**. Adjusted coefficients of determination (Adj. R^2^) for the GRF vs contact area analysis was performed using a 2nd order polynomial fit instead as this describes the data better than a linear fit. Two-way repeated measures ANOVA was used in segmentwise peak contact area analysis, as data were normally distributed according to a Shapiro-Wilk test. However, Levene’s test for homogeneity of variances was significant for SI (p<0.05) but not for ST (p=0.092), we urge caution when interpreting the within-subjects’ effects. Mauchly’s test showed sphericity of segment (W=0.082, p=0.063) and the segment*SI-ST interaction (W=0.27428, p=0.62463), where the SI-ST factor was not tested due to insufficient degrees of freedom. Independent samples t-test was performed to show no significant difference between larval behaviour on elastic resonators and standard agarose substrates as data were normally distributed according to a Shapiro-Wilk test. Pairwise comparisons between segments all used Tukey-corrected t-tests. Force-distance curves were fitted using a height-corrected Hertz Model; all force-distance curves were fitted with an R^2^>0.9.

## Acknowledgements

This work was supported by EPSRC (Doctoral Training grant EP/L505079/1 and grant EP/P030017/1), the European Research Council under the European Union’s Horizon 2020 Framework Programme (FP/2014-202) ERC grant agreement no. 640012 (ABLASE), and the Alexander von Humboldt Foundation via the Humboldt Professorship to MCG. We thank our technicians Audrey Grant and Tanya Sneddon for preparing fly food, Dr Eleni Dalaka for her support during atomic force microscopy experiments, Dr Andreas Mischok for his advice with fabrication, Dr Marcus Bischoff for his advice and support early in the work, and Dr Jacob Francis and Dr James MacLeod for their methodological advice regarding lateral view imaging.

## Author Contributions

Experiments were conceived and designed by J.H.B., M.C.G. and S.R.P.; fabrication and experiments were performed by J.H.B., method was designed by N.M.K. and A.M.; data analysis was performed by J.H.B., N.M.K. and A.M.; animal rearing was performed by J.H.B. and S.R.P; writing was performed by J.H.B. and S.R.P. with input from by M.C.G., N.M.K. and A.M.

## Supplementary Information

**Figure S1.**
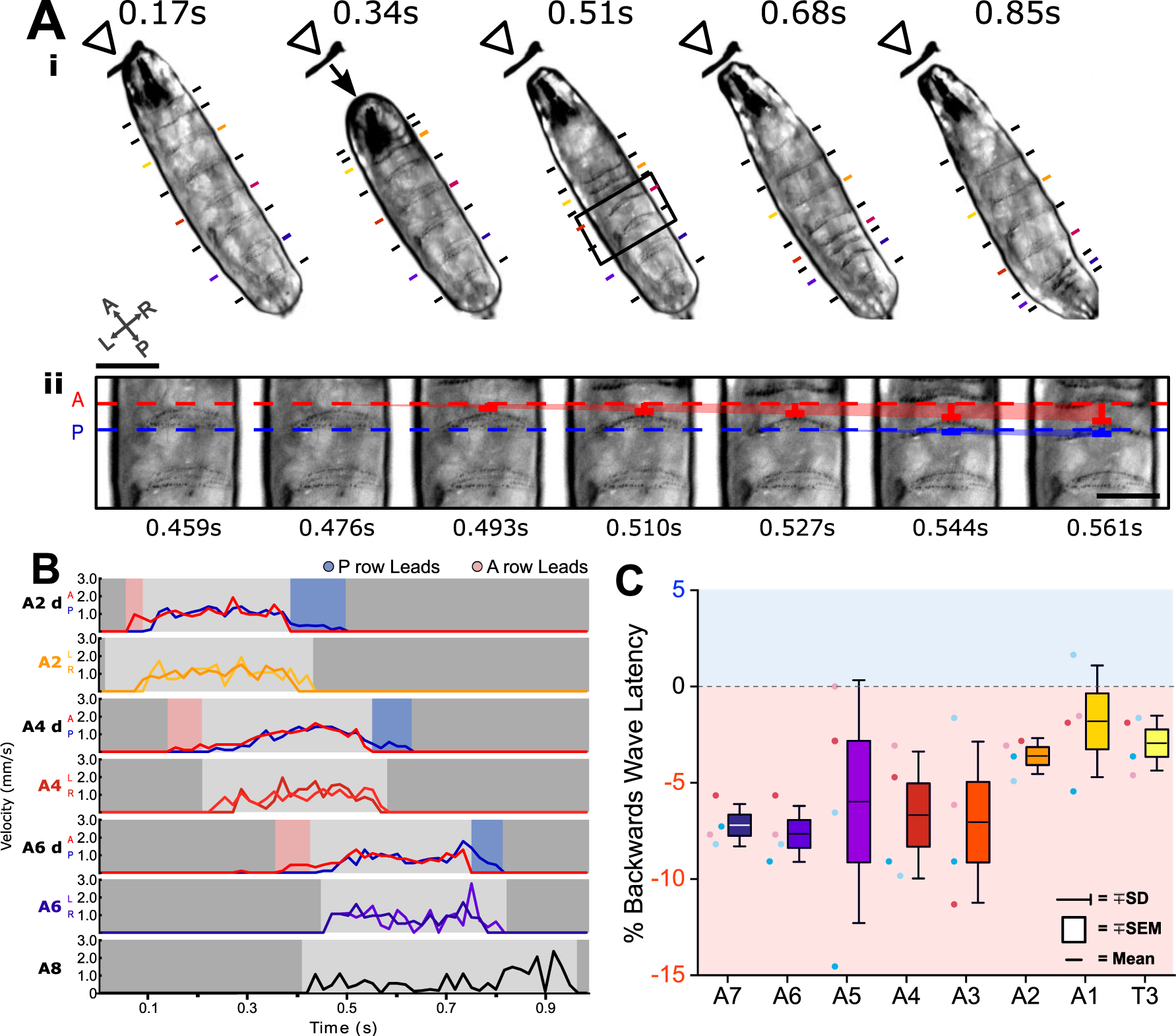
Backwards waves show a reversed heel toe rule. **A**) Backwards waves are characterised by sequential contractions moving from anterior to posterior, the reverse of forwards waves (**i**). We observed that the anterior row of each denticle band moved to meet the posterior row before the whole protopodia began to move (**ii**). **B)** We tracked the velocity of the lateral edge of each denticle band (A6, A4, A2) and the anterior and posterior rows of each denticle band (A6d, A4d, A2d). Similar to forward waves in **Figure 2**, we observed an anteroposterior latency between when each row moved relative to the other. However, this was the reverse of forwards waves, with the swing initiation period being characterised by an anterior-led latency and the swing termination period being characterised by a posterior-led latency. **C)** We found that this was relatively consistent across segments, where negative numbers represent anterior led latency, within a sample of 4 waves across 4 different animals.

**Figure S2.**
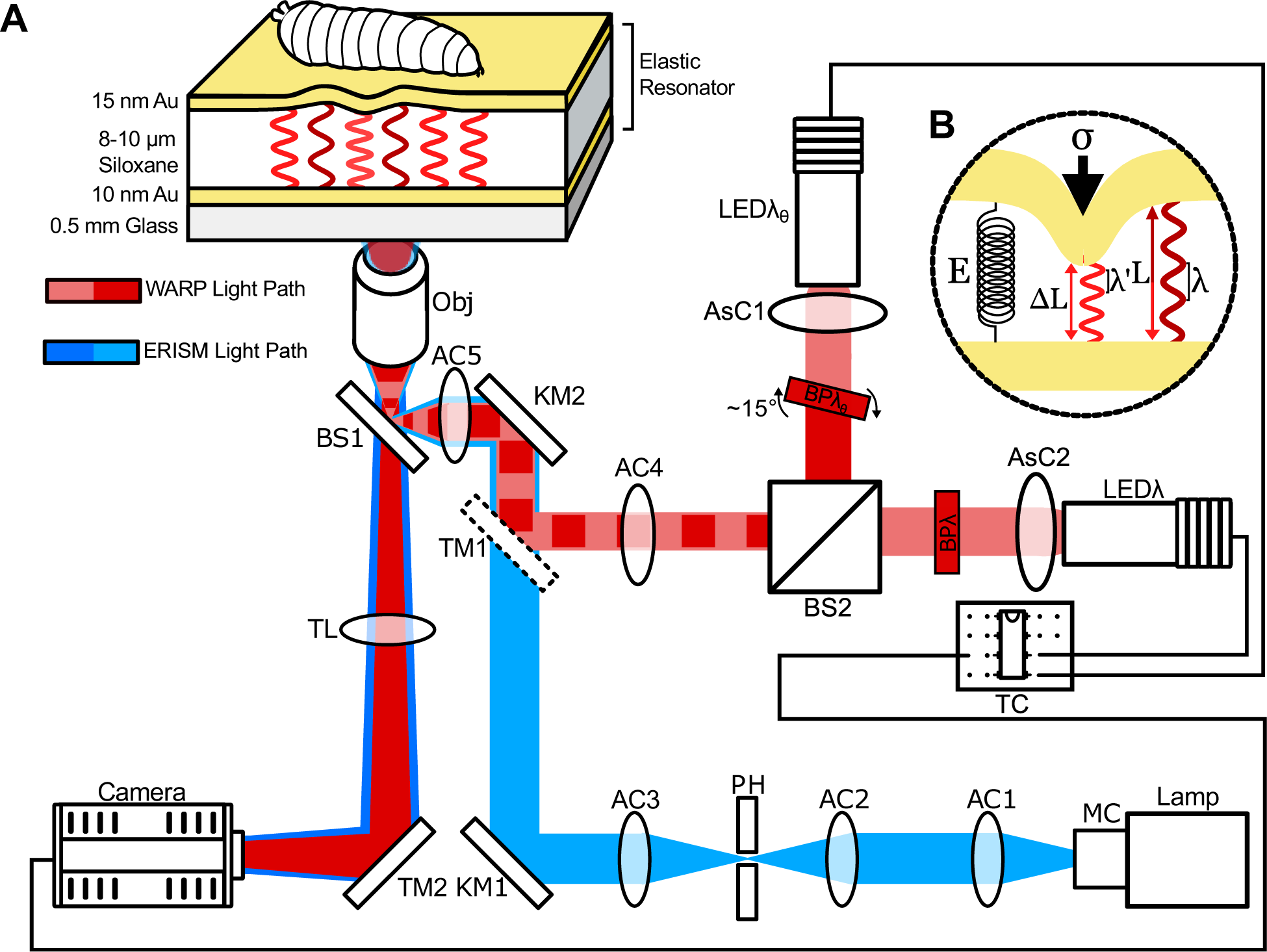
Optical setup for ERISM and WARP experiments. **A**) Optical light path used to record resonance from incident light on elastic resonators. For ERISM (blue), light originates from a halogen lamp (Lamp) and is spectrally scanned by a monochromator (MC) before being collimated by an achromatic doublet lens (AC1) and focused by another achromatic doublet lens (AC2) through a pinhole (PH). Light emerging from PH is then recollimated by AC3 into a kinematic mirror (KM1), directing it to KM2 which then directs the light under the nosewheel of a Nikon Ti2 inverted microscope. Under the nosewheel, the light is focused by an achromatic doublet lens (AC5) to a 50:50 RT beamsplitter plate (BS1), which directs the focused light to the back aperture of the objective (Obj). Light is then introduced onto the elastic resonator via Obj, whereupon it either enters the cavity, should it meet the resonance condition for the given cavity thickness, or is reflected, should it not meet this condition. Reflected light is then collected by Obj, focused by the tube lens (TL) and directed by the microscope turning mirror (TM2) then recorded by a camera (Camera). Though depicted as blue, ERISM typically scans through a spectral band from 550-750 nm. For WARP (red), light is generated by two 625 nm red LEDs (LEDλ and LEDλθ). For both LEDs, the light is collimated by aspheric condenser lenses (AsC1 and AsC2) and is then filtered by 633 nm bandpass filters (BPλ and BPλθ). BPλθ, is rotated roughly 15° such that the filter pass band is blue-shifted and the resultant transmitted light is approximately 90 degrees out of phase (in terms of the resonances of the elastic cavity) relative to light passing through BPλ. These light paths are combined by a beamsplitter cube (BS2) and collimated using AC4 into the KM2, BS1, AC5, Obj common light path by a dielectric turning mirror (TM1) only present when using WARP. The LEDs are then triggered in an alternating pattern by a trigger circuit decade counter (TC) which is controlled by the trigger out of the sCMOS camera. **B)** orking principle within the elastic cavity. hen under stress (σ), the elastic cavity deforms from its resting length (L) to its strained length (ΔL). The change in cavity length causes a change in the wavelengths that fulfil the resonance condition of the cavity. The amount of strain under a given stress is a direct consequence of the Young’s modulus (E) of the elastic material.

**Figure S3.**
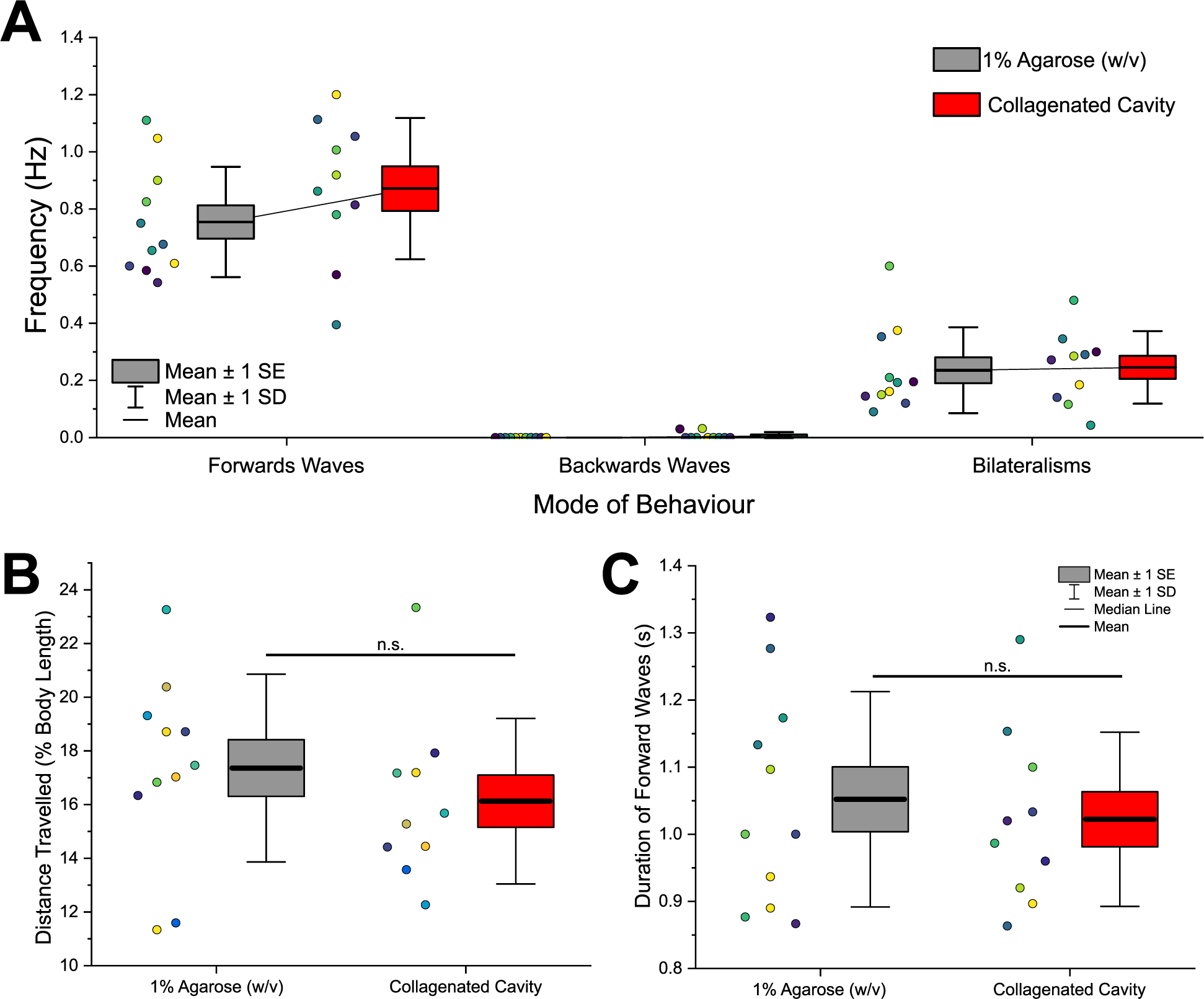
Ordinary larval behaviour is maintained on collagen treated microcavities as compared to commonly used agarose substrates. **A**) Total number of behaviours per second is not significantly different between agarose substrates and elastic cavities with collagen coating according to a two-sample t-test (t(_18_)=-1.24, p=0.23). Data for forwards waves, backwards waves and head sweep bilateralisms are shown. **B)** Distance travelled as % of body length was not significantly different between the two substrates according to a two-sample t-test (t(_18_)=1.34, p=0.20). Each data point represents the mean of 5 waves from a single animal. **C)** The mean duration of 5 forwards waves was not significantly different on microcavities compared to agarose according to a two-sample t-test (t(_18_)=0.62, p=0.54). Data taken from 10 animals. Colour of data point indicates data from an individual animal. All tested data was found to be normally distributed according to a Shapiro-Wilk test (p>0.05).

**Figure S4.**
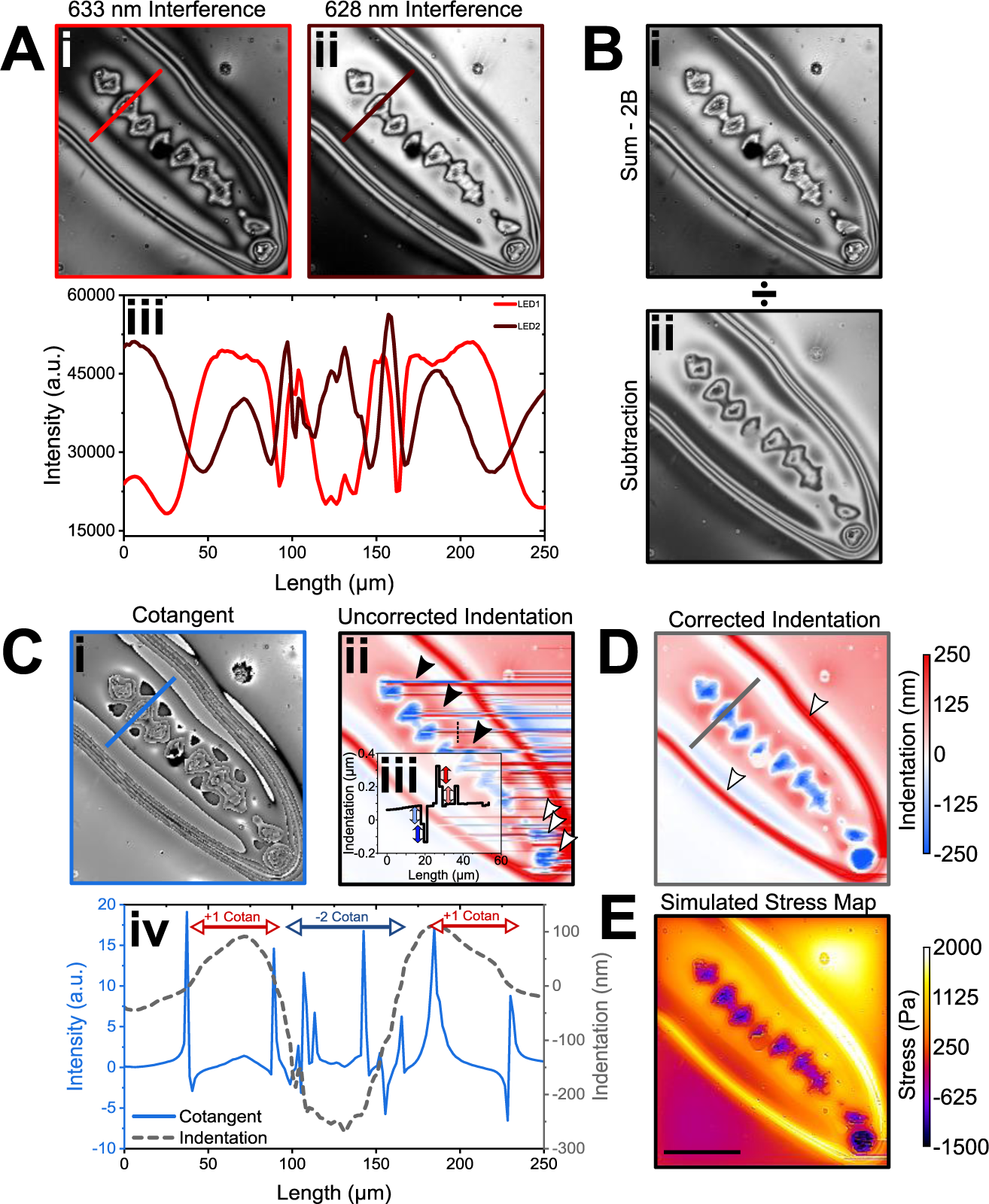
WARP computation pipeline. **A**) Interference images are taken at 633 nm (**i**) and 628 nm (**ii**) in quick alternation. Profile plot (**iii**) across the lines in **i** and **ii**. Note that interference pattern is approximately 90° out of phase relative to the other as a result of the specific wavelength difference and total cavity thickness chosen here. **B)** Images were then added together (**i**) with a background correction (2B) and then divided by the same two images subtracted from each other (**ii**). **C)** The resultant images are referred to as cotangent images as pixel intensity changes approximately as the cotangent of the cavity thickness in these (**i**). A cotangent lookup table was then used to convert 16bit greyscale values in the cotangent images to local cavity length. Raw and uncorrected displacement map computed by subtracting a linear plane of mean cavity thickness (**ii**). Profile plot (**iii**) along the thin dashed line in ii, clearly showing discontinuity artefacts (indicated by blue and red double arrows). These linear artefacts correspond to step heights amounting to jumps by one free spectral range, which was 112 nm in this instance. **D)** Corrected displacement map where the linear artefacts across the image were corrected by applying a continuity condition. **E)** Profile plots of cotangent signal and corrected displacement along the thick lines in **Ci** and **D**. **F)** Stress maps were calculated from the displacement map by FEM using the known mechanical properties of the substrate. Scalebar = 200 µm.

**Figure S5.**
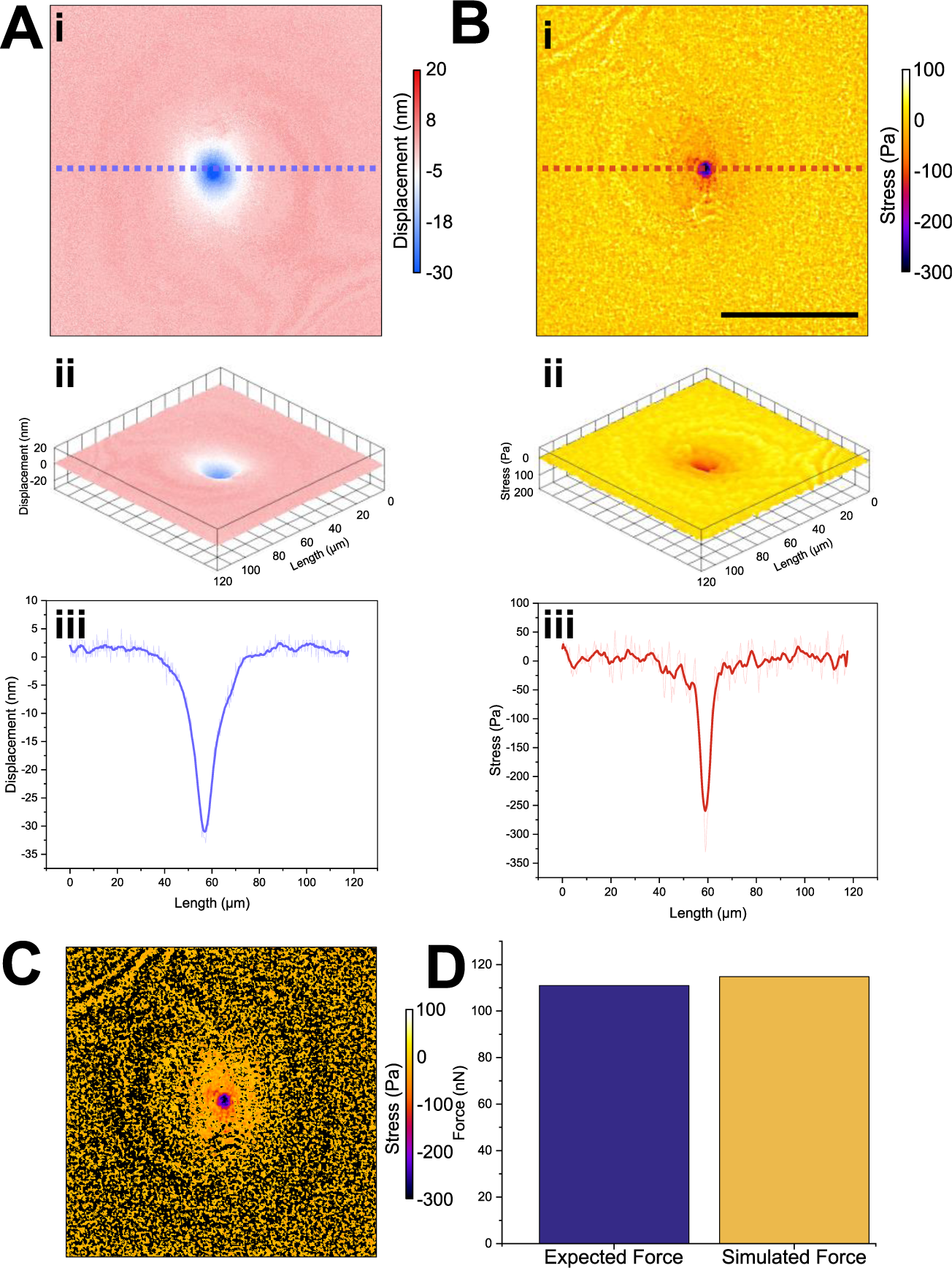
Confirmation of finite element method (FEM) simulation accuracy. **A**) Using atomic force microscopy (AFM), we indented into an elastic resonator made within the same batch as those used for videorate stress mapping. (**i**) False colour map and (**ii**) 3D projection of displacement smoothed by 10 points. (**iii**) Profile along blue dotted line in (**i**) showed that a 111 nN indentation force resulted in a roughly 32 nm peak indentation; data averaged by 10 points, raw data shown in a lighter colour. **B)** Using FEM, we calculated a stress map from the displacement map, using the young’s modulus of the bulk material, previously recorded as 16450 Pa by AFM. (**i**) False colour and (**ii**) 3D projection of stress experienced by the resonator smoothed by 10 points (**ii**). Profile along red dotted line in (**i**) showing the peak stress produced by 111 nN of force approximately 320 Pa; data averaged by 10 points, raw data shown in a lighter colour. **C)** We thresholded the resultant simulation to remove all cavity displacements >0nm. **D)** Integration of stress in **C** gives a prediction of the total applied force as determined from displacement map and FEM model, without prior knowledge of the indentation force. Comparing this simulated force to the applied force of 111 nN, we found the relative difference to be only 3.4%, with our simulation estimating a total applied force of 114.8 nN. Scale bar in **B** denotes 50 µm.

**Table S1:** Measured effective Young’s Modulus per elastomer mixture post plasma treatment.

| **Elastomer Mixture** | **Mean Young's Modulus (Pa)** | **Plasma power (W)** |
| --- | --- | --- |
| 10% Sylgard®184,<br>90% Sylgard®527 | 27697.978 ± 913.45 | 30 |
| 6.6% Sylgard®184,<br>94.4% Sylgard®527 | 25999.17 ± 991.93 | 30 |
| 5% Sylgard®184,<br>95% Sylgard®527 | 18596.08 ± 186.05 | 30 |
| 100% Sylgard®527 | 12325.35 ± 395.45 | 30 |
| 100% Nusil®GEL8100 | 3340.30 ± 53.45 | 30 |
| 10% Sylgard®527,<br>90% Nusil®GEL8100 | 8974.32 ± 621.83 | 180 |
| 50% Sylgard®527,<br>50% Nusil®GEL8100 | 18404.14 ± 439.85 | 180 |
| 90% Sylgard®527,<br>10% Nusil®GEL8100 | 38387.87 ± 2768.63 | 180 |

**Video SV1.**
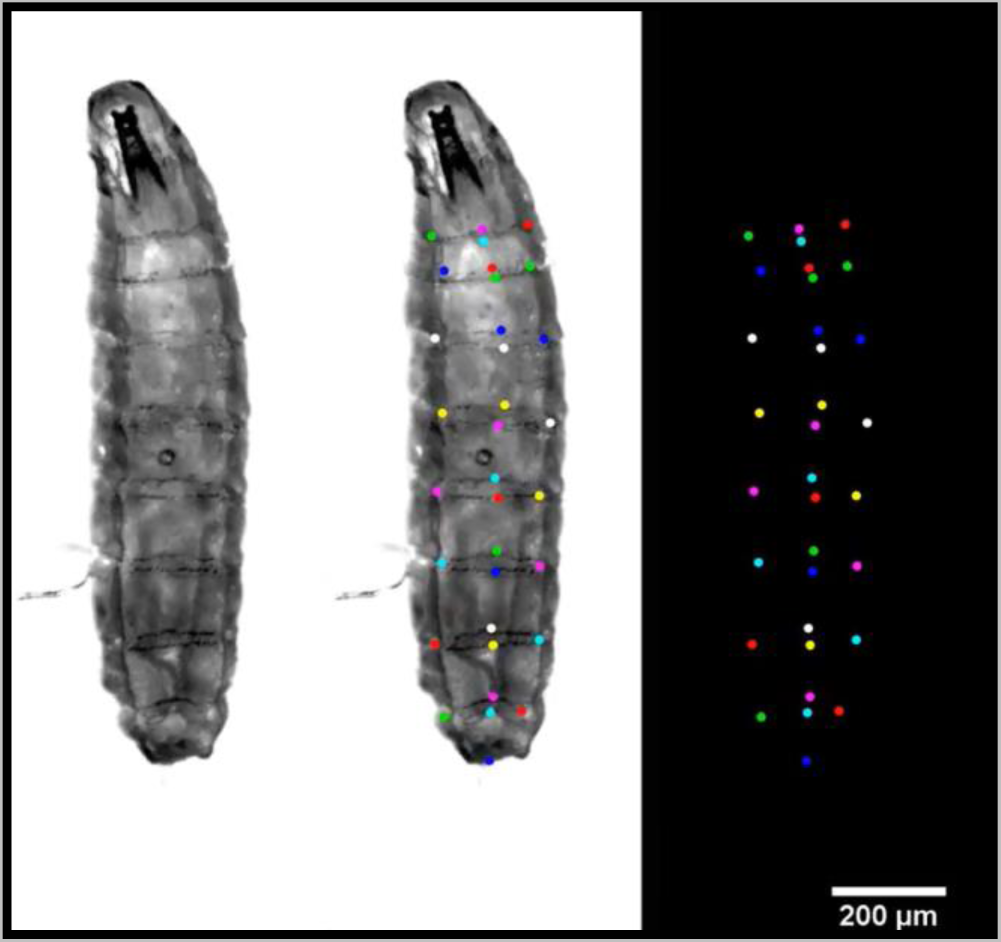
Kinematic Tracking of Forwards and Backwards Peristaltic waves. Manual tracking of 33 points across the body during forwards and backwards peristalses.

**Video SV2.**
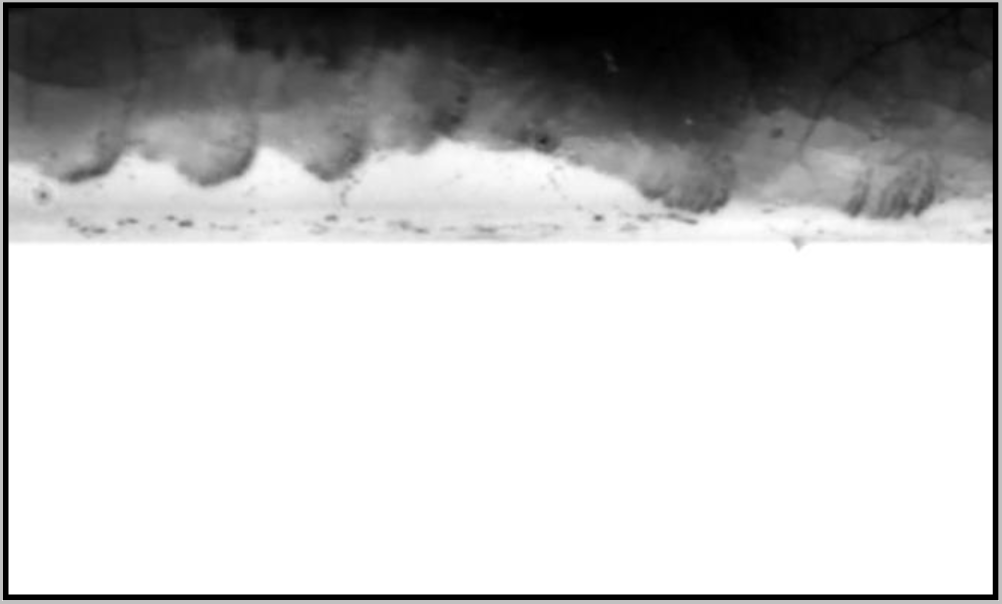
Lateral view crawling. Video showing the sequestration and planting of protopodia during locomotion from a lateral view.

**Video SV3.**
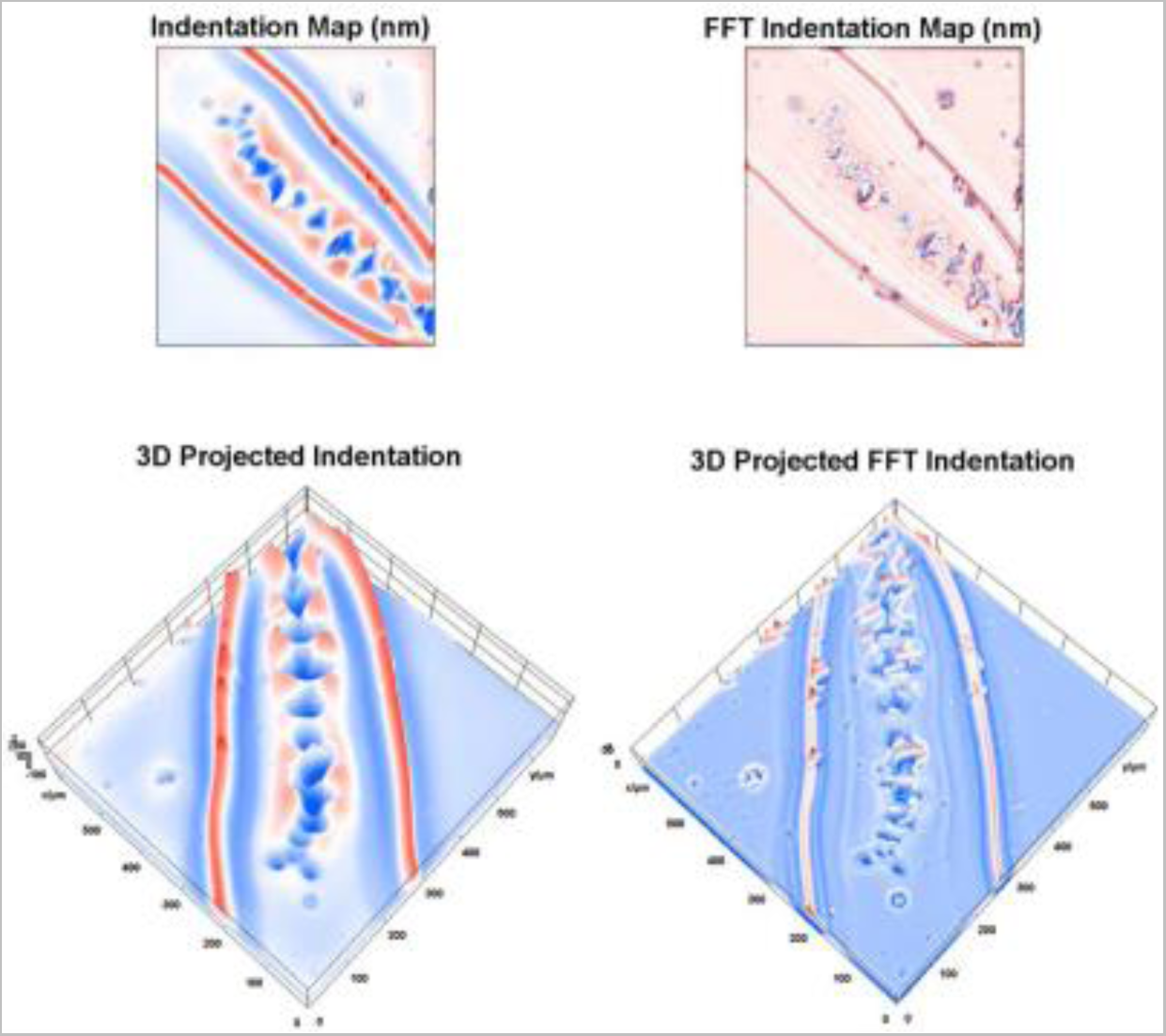
WARP imaging during forwards peristalses. Video showing high frame rate displacement maps produced by a freely behaving Drosophila larva. Displacement maps were high-pass Fourier filtered to make denticulated cuticle more readily visible and projected in 3D to show the effects of substrate interaction. Details of the Fourier filtering procedure were described in a previous study (23).

**Video SV4.**
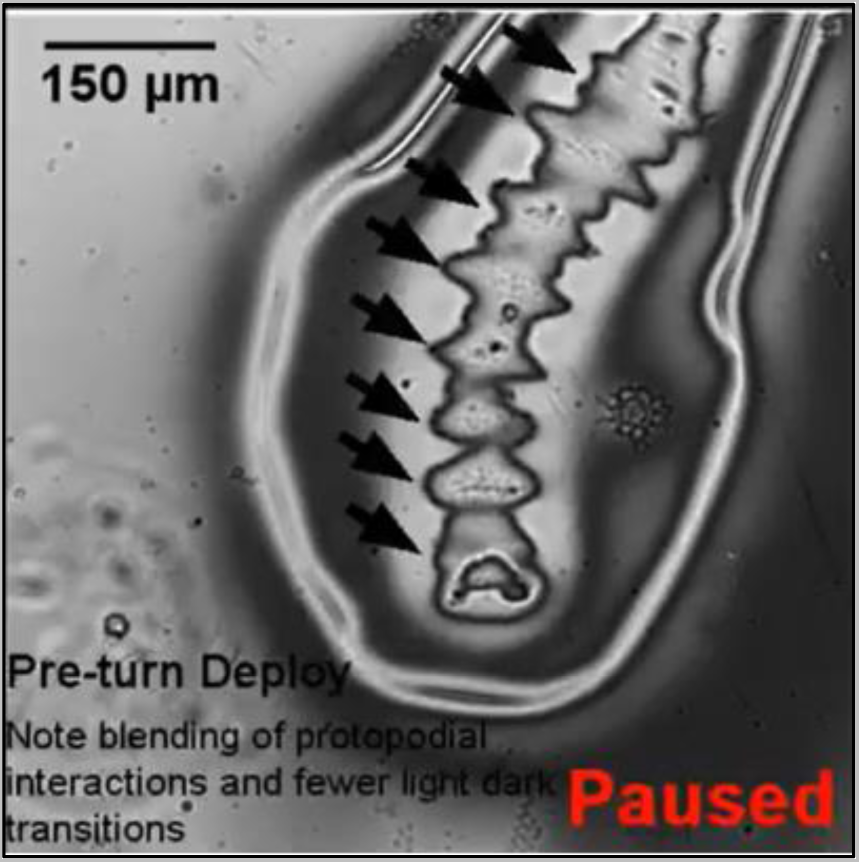
Interference mapping of bodymass redistribution during anterior bilateral behaviours. Video showing the raw reflection data during the preparatory phase of bilateral behaviours.

